# Repeated translocation of a supergene underlying rapid sex chromosome turnover in *Takifugu* fish

**DOI:** 10.1101/2021.11.16.468883

**Authors:** Ahammad Kabir, Risa Ieda, Sho Hosoya, Daigaku Fujikawa, Kazufumi Atsumi, Shota Tajima, Aoi Nozawa, Shotaro Hirase, Takashi Koyama, Osamu Nakamura, Mitsutaka Kadota, Osamu Nishimura, Shigehiro Kuraku, Yasukazu Nakamura, Hisato Kobayashi, Atsushi Toyota, Satoshi Tasumi, Kiyoshi Kikuchi

## Abstract

Recent studies have revealed a surprising diversity of sex chromosomes in vertebrates. However, the detailed mechanism of their turnover is still elusive. To understand this process, it is necessary to compare closely related species in terms of sex-determining genes and the chromosomes harboring them. Here, we explored the genus *Takifugu*, in which one strong candidate sex-determining gene, *Amhr2*, has been identified. To trace the processes involved in transitions in the sex determination system in this genus, we studied 12 species and found that while the *Amhr2* locus likely determines sex in the majority of *Takifugu* species, three species have acquired sex-determining loci at different chromosomal locations. Nevertheless, the generation of genome assemblies for the three species revealed that they share a portion of the male-specific supergene that contains a candidate sex-determining gene, *GsdfY*, along with genes that potentially play a role in male fitness. The shared supergene span approximately 100 kb and are flanked by two duplicated regions characterized by CACTA transposable elements. These results suggest that the shared supergene has taken over the role of sex-determining locus from *Amhr2* in lineages leading to the three species, and repeated translocations of the supergene underlie the turnover of sex chromosomes in these lineages. These findings highlight the underestimated role of a mobile supergene in the turnover of sex chromosomes in vertebrates.

**Significance:** Although turnover of sex chromosomes is very common in many vertebrate lineages, the transition process is still elusive. We studied the sex-determining region (SDR) of 12 congeneric fish species. We found that while nine species retained their ancestral SDR, three species had acquired derived SDRs. Although the derived SDRs resided in three different chromosomes, they harbored a shared supergene flanked by two putative transposable elements. The results highlight the underestimated role of a mobile supergene in turnover of sex chromosomes in vertebrates.

## Introduction

Sex-determining genes and the chromosomes harboring them have been maintained in therian mammals and birds for more than 100 million years (1). However, such stability and conservation of the sex determination system are not universal in vertebrates. Indeed, it has been shown that sex chromosomes have changed (“turned over”) in many poikilothermic vertebrate lineages, such as fishes, amphibians, and non-avian reptiles (2–7).

Among them, fishes are a particularly attractive group of animals in which to investigate the rapid turnover of sex chromosomes and/or sex-determining genes, since different sex determination mechanisms often exist within a group of closely related species, and the sex-determining genes have been identified in some of these groups. However, in the majority of cases, distinct sex-determining genes were only identified in phylogenetically distant species.

Therefore, it is challenging to illuminate key genomic changes associated with the evolutionary transition between the ancestral and derived sex-determination systems, and hence evolutionary drivers facilitating the transition remain largely vague. Although extensive comparative studies have been ongoing in some genera, each of which includes a relatively well-studied species such as medaka, guppy, pejerrey, Nile tilapia, threespine stickleback, rainbow trout, and northern pike (8–17), the transition between the ancestral and derived sex-determining genes have only been traced at the molecular level in medaka and its relatively distant species that diverged more than 20 million years ago (18, 19).

*Takifugu* is a genus of fish that includes fugu (*T. rubripes*), a model species for comparative genomics (20), and approximately 20 closely related species (21) (Fig. 1A). It has been known that sex chromosomes in the lineage leading to fugu have remained undifferentiated for more than 4 million years (1, 15). The anti-Müllerian hormone receptor type II (*Amhr2*) is most likely the sex-determining gene in fugu, *T. pardalis*, and *T. poecilonotus*, since the single nucleotide polymorphism (SNP; SNP7271) on exon9 of the gene was the sole polymorphism associated with phenotypic sex (G/C in male) and there is no sign of sequence differentiation beyond this SNP between the X and Y chromosomes (15, 16). However, a recent study of this genus showed a possible transition of the sex-determining genes in one of the congeneric species, *T. niphobles*, in which the sex-determining locus mapped to a location distinct from that of *Amhr2* (15, 16). Moreover, the strong suppression of recombination around the sex-determining locus in *T. niphobles* suggested that this species potentially has sex chromosomes with an early stage of sequence differentiation. Since the ancestor of the *Takifugu* genus underwent rapid speciation 2–5 million years ago, the genus will provide a unique opportunity for understanding the process of rapid sex chromosome turnover and the initiation of sex chromosome differentiation, as well as the genomic changes associated with each of these.

**Fig. 1.**
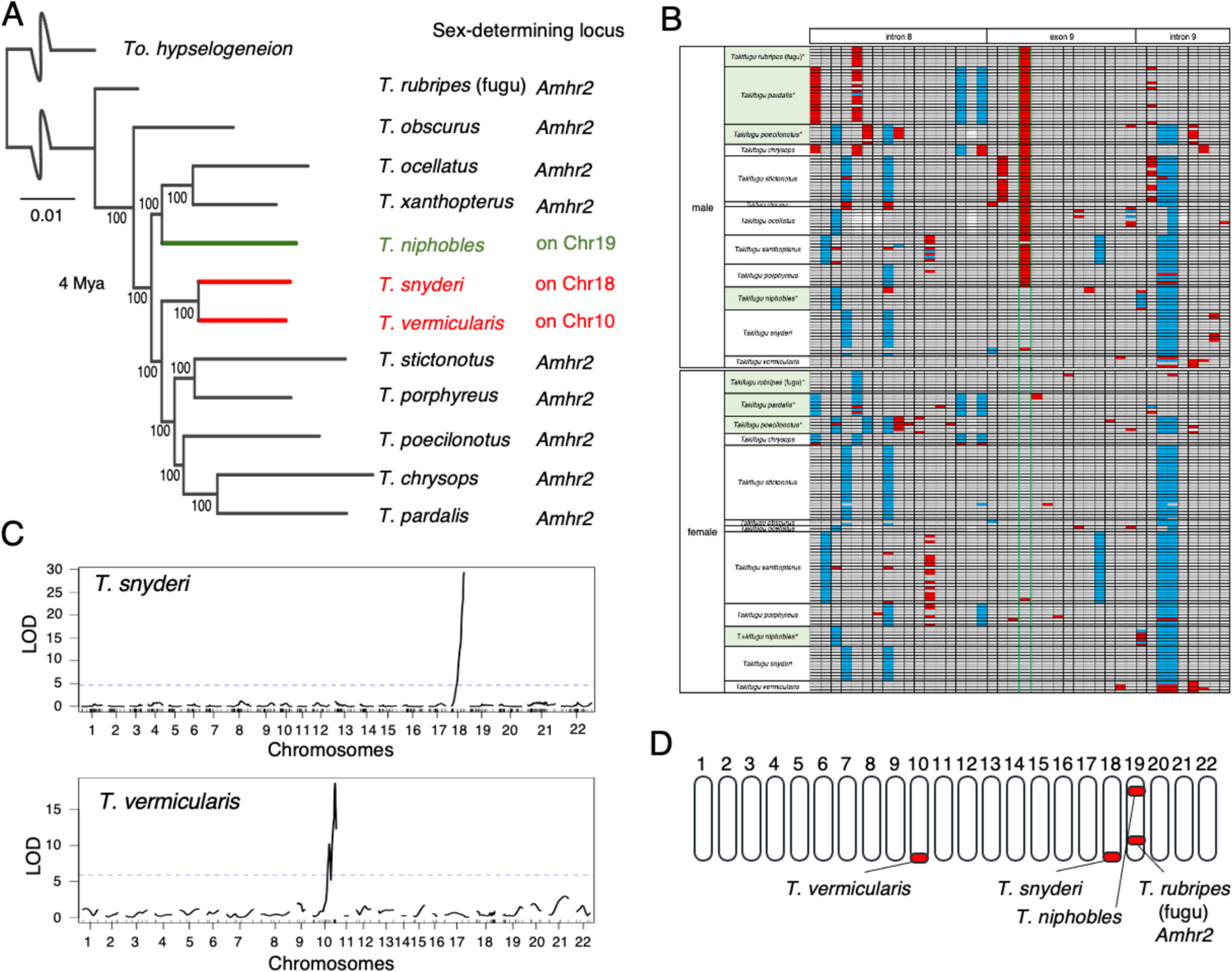
The genus *Takifugu* contains at least three species with sex chromosome turnover. (A) Phylogenetic relationship of 12 *Takifugu* species and their sex-determining loci. The green and red lines indicate lineages with a distinct sex-determining locus from *Amhr2*. Bootstrap values are shown at the nodes. Trees were rooted with *Torquigener hypselogeneion* as an outgroup. Divergence times (4.22 MYA, CI: 2.58-6.21 MYA) at the crown age of the genus are based on fossil records and mitochondrial and transcriptome divergence (98). (B) A comparison of SNPs in *Amhr2* among 12 *Takifugu* species suggests that sex is not determined by the SNP7271 in the three species with sex chromosome turnover. *The data for four fugu species (*T. rubripes* (fugu), *T. pardalis*, *T. poecilonotus*, and *T. niphobles*) are taken from ref. (16) for comparison (shaded in pale green). SNP sites detected in the partial sequence of *Amhr2* (intron 8, exon 9, and intron 9) are shown. SNP7271 of *Amhr2* is flanked by green lines. Gray cells indicate that an individual is homozygous for the reference allele, blue cells indicate that an individual is homozygous for the alternative alleles, and red cells indicate that an individual is heterozygous. White cells refer to deletions. Source data underlying this panel are provided as a Source Data file. (C) Linkage analyses map the sex-determining locus to Chr18 in *T. snyderi* and to Chr10 in *T. vermicularis*. QTL analysis was conducted using 1,225 markers for 98 full-sib progenies in *T. snyderi* Family A, and using 139 markers for 62 full-sib progenies in *T. vermiculari*s Family D. The LOD score is on the Y-axis, and linkage groups (chromosomes) are on the X-axis. Chromosome identity is based on conserved synteny with the reference genome of fugu (*T. rubripes*) defined by ref. (23) (FUGU5/fr3, GenBank: GCA_000180615.2). The marker position is indicated on the X-axis. The significance level (p = 0.05) is indicated by dashed lines. Source data underlying these panels are provided as a Source Data file.

To gain insight into the transition process between the sex determination systems, we expanded our previous work to include 12 *Takifugu* species and found that sex chromosome replacements have occurred in three species. We then generated a genome assembly for each of the three species and unexpectedly found that despite the fact that the chromosomal location of the sex-determining locus is distinct among the three species, these loci share a very similar male-specific region. We speculated that repeated translocation of a pre-existing sex-determining locus may be more prevalent in vertebrates than previously thought, and may have caused rapid turnover of sex chromosomes.

## Results

### The sex-determining SNP in *Amhr2* is absent in three of 12 species

Previous studies suggested that the sex-determining role of SNP7271 in *Amhr2* is not conserved in one of four *Takifugu* species, namely *T. niphobles* (16) (shown in green in Fig. 1A). To gain a more comprehensive view of the transition of sex-determining loci in this genus, we sequenced the genomic segment harboring the SNP7271 site from females and males of eight *Takifugu* species that had not been examined previously (Fig. 1B, Table S1). We found that while the association between SNP7271 polymorphisms and gonadal sex was conserved among six species, this was not the case in the other two; the SNP locus was homozygous for the C allele in all individuals of *T. snyderi* and *T. vermicularis*, regardless of their sex (Fig. 1B, Table S1, shown in red in Fig. 1A). We further confirmed the absence of the G allele at the SNP site in wild populations of the two species (n = 50 for each sex of each species) by determining the genotype using a high-resolution melting assay (22) (Table S1).

**Table 1.** Assembly statistics for the genomes of T. niphobles, T. snyderi, and T. vermicularis.

|  | *T. niphobles* | *T. snyderi* | *T. vermicularis* |
| --- | --- | --- | --- |
| Genotype | YY | XY | XY |
| Sequencing and scaffolding strategies | PacBio, Illumina, Hi-C | Nanopore, stLF, Illumina | PacBio, Illumina |
| The number of scaffolds/contigs | 424 scaffolds | 1,097 scaffolds | 1,756 contigs |
| Total length | 396 Mb | 399 Mb | 429 Mb |
| Average length | 935 kb | 363 kb | 244 kb |
| Longest scaffold/contig | 29.8 Mb | 22.3 Mb | 8,669 kb |
| Shortest scaffold/contig | 1,000 bp | 162 bp | 2,979 bp |
| Scaffold/contig N50 | 16 Mb | 6 Mb | 1.6 Mb |

### Linkage mapping identifies novel sex-determining loci

The genotyping results implied that the sex-determining loci have turned over in lineages leading to *T. snyderi* and *T. vermicularis*. Alternatively, the role of master sex determiner in these species may have been taken over by environmental factors. To test these hypotheses, we produced 98 full-sib progenies (Family A) in *T. snyderi* and 62 full-sib progenies (Family D) in *T. vermicularis* from wild-caught parents (Table S2), and performed genome-wide linkage mapping of the sex-determining locus. We found a strong linkage between gonadal sex and paternally inherited alleles at loci near the distal end of Chr18 in *T. snyderi* (LOD = 29.3, *P* = 1.8 × 10^-23^) and Chr10 in *T. vermicularis* (LOD = 18.7, *P* = 4.3 × 10^-15^) (Fig. 1C). We further confirmed this result in additional families of *T. snyderi* (*P* < 1.0 × 10^-16^) and *T. vermicularis* (*P* = 1.8 × 10^-39^) by marker association (Fisher’s exact test) (Table S3). The linkage analysis also revealed that the marker loci and the synteny of these markers are largely conserved between the species (Fig. S1).

### A phylogenetic framework suggests three independent turnovers

Summing up the above results and those of previous studies (15, 16), it is now evident that while the role of the *Amhr2* locus in sex determination is largely conserved among *Takifugu* species, at least three turnover events have occurred in this genus, resulting in four distinct sex-determining loci (Fig. 1D). To infer the ancestral and derived states of the four sex-determining loci, we generated a phylogenetic tree using whole-genome resequencing data of 12 *Takifugu* species and one outgroup species (Fig. 1A, Table S4). According to the most parsimonious interpretation of the phylogeny, the results corroborate the previous hypothesis that *T. niphobles* has obtained a derived sex-determining locus on Chr19, while having lost the ancestral male-determining allele (the G allele) at the *Amhr2* locus on the same chromosome (16) (green in Fig. 1A). Moreover, the phylogenetic tree indicates that sex chromosomes are likely to have turned over at least twice in the lineage leading to the sister species, *T. snyderi* and *T. vermicularis* (red in Fig. 1A).

Having detected the three turnover events related to sex chromosomes within *Takifugu* species, we turned to the detailed characterization of the three sex-determining loci that have likely arisen recently (shown in green and red in Fig. 1A).

### *k*-mer analysis identifies male-specific sequences in *T. niphobles*

Although we previously showed that the sex-determining locus in *T. niphobles* maps to Chr19 with an XX/XY system (16) (shown in green in Fig. 1A), no male-specific sequences have been found. To identify male-specific sequences in *T. niphobles*, we first searched for the male-specific *k*-mers in the resequencing data of six females and five males (Table S5) and obtained the 35-mer sequences unique to males. Using these sequences, we extracted the pair-end reads from the same resequencing data and assembled them into 551 contigs with a total of 218-kb sequences (Table S6). A BLAST search of the contigs against a fugu reference genome sequence, FUGU5 (23), revealed that similar sequences exist on multiple chromosomes in *T. rubripes*, including Chr4, Chr6, and Chr7 (Table S7). The result raised the possibility that the male-specific region on the Y chromosome (Chr19) in *T. niphobles* may have been derived from multiple autosomal regions.

### Construction of a chromosomal-scale genome assembly of a *T. niphobles* YY male

As the *k*-mer approach suggested that the male-specific region of *T. niphobles* is likely absent in the fugu reference genome, we attempted to generate a reference genome sequence for *T. niphobles*, including its Y chromosome and the male-specific region. Complete sequence assemblies of Y chromosomes have not been generated for many organisms (but see some exceptions in (24–28)) because of technical difficulties related to the discrimination of X and Y in the male genome and/or the assembly of regions containing high levels of repetitive sequences.

To alleviate these problems, we sequenced the genome of a male *T. niphobles* carrying two Y chromosomes (instead of X and Y) and generated a draft genome sequence using a hybrid assembly strategy with PacBio long reads and Illumina short reads (Fig. S2, Table S8 and S9). The assembly consists of 728 contigs with an N50 of 6.08 Mb and a total size of ∼396 Mb.

Further scaffolding with Hi-C data placed 288 contigs into 22 superscaffolds (pseudochromosomes) that had chromosomal-level length (longer than 10 Mb) (Fig. S3B). The size of these pseudochromosomes covered 91.2% of the total genome and their number was consistent with the karyotype of this species (29) (Fig. S3B, Table S10 and S11). The total number of scaffolds in the Hi-C–based proximity-guided assembly was 424. The scaffolding increased N50 from 6.1 Mb to 16.2 Mb and that of the longest scaffold from 15 Mb to 29.8 Mb (Table 1, Fig. S3B). A BUSCO search against the 4,584 single-copy orthologs for Actinopterygii suggested that the assembly was missing only 2.3% of the core genes (Table S12). Furthermore, approximately 95% of the short reads from multiple male individuals were uniquely mapped to the assembled genome with a mapping quality above 20. These metrics indicated the high quality of the chromosomal scale assembly. The assembly showed highly conserved synteny with the fugu reference genome (Fig. S4), and therefore chromosome identities for the *T. niphobles* were assigned based on those of *T. rubripes*. Based on the conserved synteny shown here and the genomic positions of the sex-linked markers reported previously (16), we identified the pseudochromosome corresponding to the *T. niphobles* sex chromosome (Chr19), which comprises 23 contigs totaling 17 Mb in length (Fig. S5, Table S11). Because this pseudochromosome was assembled from the YY male, we denoted it as Chr19Y.

### The male-specific region is flanked by pseudoautosomal regions in *T. niphobles*

To determine the male-specific region in this assembly, we aligned resequencing reads from 8 females and 10 males against the assembly and compared the relative depth of coverage between males and females. We detected a clear difference in the coverage in the region (∼245-kb) on Chr19Y (Fig. 2A), suggesting that the region may be specific to males. The male-specific region is composed of two sub-regions with different relative coverage; the left-half region (130-kb) showed a relatively higher male-to-female ratio than the right-half region (115-kb) (Fig. 2A). The absence of differences in coverage depth between sexes along the flanking regions indicates that these regions are shared between Y and X chromosomes or have not diverged enough to be distinguished by uniquely mapped reads, thus delineating the boundaries of the male-specific region and pseudoautosomal regions.

**Fig. 2.**
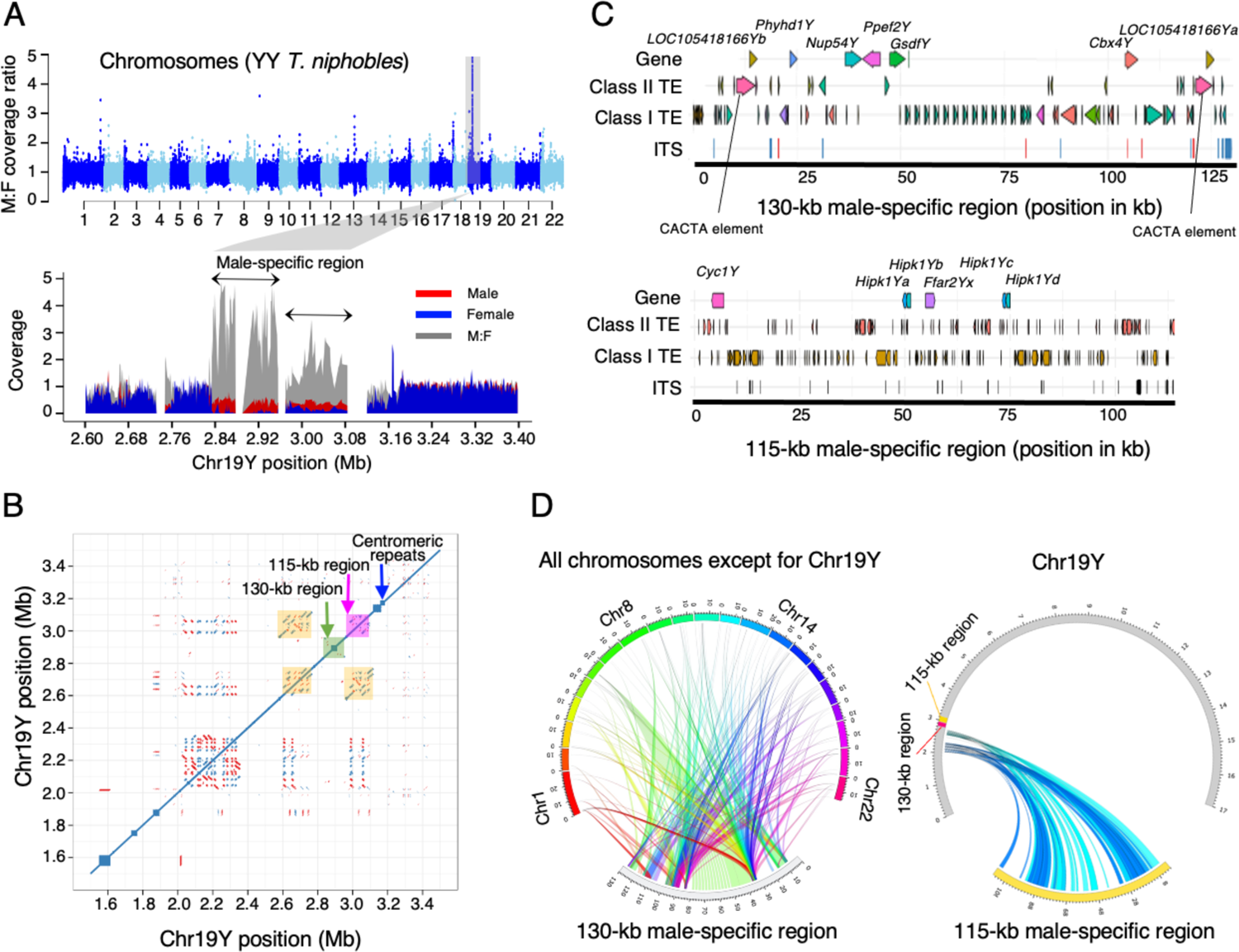
The male-specific region in *Takifugu niphobles*. (A) The male-specific region in *T. niphobles* was revealed by differences in the depth of coverage between males and females. We aligned resequencing reads from 10 males and 8 females against the assembled genome of a *T. niphobles* YY male and calculated the relative depth of coverage between sexes ((average male coverage+1) / (average female coverage+1)) at a window size of 1-kb (Y-axis). In the upper plot for the genome-wide analysis, chromosomes are on the X-axis. In the lower plot, the male-specific region and part of the pseudoautosomal region on Chr19Y are shown. The normalized coverage of males, females, and their ratio are shown in red, blue, and gray, respectively. (B) A self-comparison of the Y-chromosome sequence suggests the accumulation of large repetitive sequences in half of the male-specific region. The dot plot represents a self-comparison of the genomic region in the vicinity of the male-specific region (the positions from 1.5 to 3.5 Mb). The left half (130 kb) and right half (115 kb) of this region are shaded in green and dark pink, respectively. Marked sharing of the repetitive sequence between the right-half region and a pseudoautosomal region is shaded in orange. Direct (blue) and inverted (red) repeats are visualized as the accumulation of dots off the central diagonal line. The blue arrow indicates the accumulation of a centromeric repeat. (C) Schematic representations of the repeat annotation in the male-specific region in *T. niphobles*. Each haplotype is annotated with triangles to depict annotated full-length genes, Class I transposable elements (TEs), and Class II TEs. Rectangles represent interstitial telomeric sequences (ITSs) (TTAGGG)n>=2. (D) Comparison of the male-specific region and other parts of the genome in *T. niphobles*. In the left plot, the 130-kb male-specific region is scaled in kb and the chromosomes are scaled in Mb. The results suggest that autosome-to-Y transposition is the main source of the formation and expansion of the male-specific region. In the right plot, Chr19Y is scaled in Mb. The results suggest that the 115-kb region originated from segmental duplications on a pseudoautosomal region.

### Accumulation of repetitive sequences is heterogeneous in the male-specific region of *T. niphobles*

Accumulations of large repetitive sequences have been observed in the non-recombining regions of sex chromosomes across diverse taxa (30, 31). To determine if this is the case in *T. niphobles*, we aligned Chr19Y to itself and found that the repeat accumulation was still modest in about half of the male-specific region (130 kb) (Fig. 2B; shaded in green), while the other 115-kb contained many large repetitive sequences (Fig. 2B; shaded in pink). Furthermore, the self-alignment plot implied that the 115-kb region is likely derived from the duplication of a region residing on the pseudoautosomal region near the male-specific 130-kb region (Fig. 2B; shaded in yellow).

### Repetitive sequences are accumulated in the male-specific region of *T. niphobles*

The repeat sequence analysis revealed accumulation of repeat sequences within the 245-kb male-specific regions; repeat sequences occupied 42.5% and 62.2% of the 130-kb and 115-kb regions, respectively (Fig. 2C, Table S13). While these values are greater than that of the whole Chr19Y (14.9%), the pseudoautosomal regions near the male-specific region also harbor repeat-rich sequences (Fig. S5). For example, repeat sequences occupied 46.3% of the ∼1-Mb pseudoautosomal region (ranging from the position 2 Mb to the proximal boundary to the male-specific region).

### A candidate sex-determining gene in *T. niphobles* resides in the male-specific region flanked by putative transposable elements

Since the *k*-mer analysis suggested that part of the male-specific region may have been derived from an autosomal region, we tested this possibility by performing BLAST searches of the male-specific sequences (130 kb + 115 kb) against the reference genome sequences of *T. niphobles* and *T. rubripes* (FUGU5). We found that the majority of the 130-kb male-specific region was composed of small segments that showed high sequence similarity to portions of other chromosomes, suggesting that the transposed sequences were the main source of the male-specific region (Fig. 2D). Furthermore, we annotated seven protein-coding genes that are paralogous to the genes on three different autosomes (Fig. 2C): *GsdfY*, *Ppef2Y*, *Nup54Y*, *Phyhd1*, *Cbx4Y*, and two *LOC105418166Y* (uncharacterized protein-coding genes that appear to be conserved in teleost fish). Of these, *GsdfY* is worth noting since the ortholog of this gene has been reported as the master sex-determining gene in a teleost species, *Oryzias luzonensis* (18). Furthermore, loss of function of this gene was found to result in female-to-male sex reversal in two teleost species, Nile tilapia and medaka (34, 35). Our RNA-seq analysis confirmed the paralog-specific expression from this locus in differentiating male gonads, along with the expressions of *Ppef2Y* and *Nup54Y* (Fig. S6, Table S14). As for *Phyhd1Y*, *Cbx4Y*, and *LOC105418166Y*s, their expression of either the male-specific or autosomal paralogs was observed but paralog-specific expression could not be ensured due to the lack of diagnostic nucleotide sites that can distinguish the transcripts of these paralogs.

Intriguingly, the sequence annotation also revealed that the male-specific region is flanked at both ends by two CACTA transposable elements containing the catalytic center of the transposase (Fig. 2C). CACTA elements, also described as EnSpm elements, belong to the Class II transposons that utilize a cut-and-paste mechanism for their transposition (32–34). A full-length CACTA element contains two open reading frames, one encoding a transposase and the other a protein of unknown function. Although we were unable to show solid evidence that the two CACTA elements of *T. niphobles* are structurally intact, we found that the catalytic center of the transposase called the DDD/E domain is conserved (Fig. S7).

The search involving the other half of the male-specific region (115 kb) identified six protein-coding genes (*Cyc1Y*, *Hipk1Ya*, *Hipk1Yb*, *Hipk1Yc, Hipk1Yd* and *Ffar2Y*), though their roles in sex determination have not been reported (Fig. 2C). Each gene had more than one untruncated or truncated paralog on Chr19Y at the outside of the male-specific region (Fig. 2D), corroborating the hypothesis that the 115-kb region originated from duplication events on the pseudoautosomal regions (Fig. 2B). Among these paralogs, the expression of either the male-specific or pseudoautosomal paralogs of *Cyc1Y* was observed in the developing gonads. However, the paralog-specific expression was not assessed due to the lack of diagnostic nucleotide sites.

### Verification of the male-specific region of *T. niphobles*

To verify the overall accuracy of the assembled sequences of the 130-kb male-specific region and its adjacent pseudoautosomal regions, we designed 12- and 5-primer pairs targeting each of the two regions, respectively (Table S15), and successfully obtained all amplicons. Sequence alignment revealed that the amplicon contigs were largely concordant with the target region of the Chr19Y assembly (Fig. S8). Discordant regions comprised only 1.2% of the amplicon contigs and 4.5% of the target region of the Chr19Y assembly. The mismatch may have been due to misassembly, misamplification of amplicons, or true polymorphisms between individuals used for the tiling PCR and the Chr19Y construction. We also confirmed the male specificity of the region using a wild population (n = 20 for each sex, P = 3.0 × 10^-10^) with a diagnostic primer pair that could yield amplicons of different size from the male-specific region (352 bp) and its paralogous region (408 bp) (Fig. S9). Although one potential sex-reversed fish was observed, similar incomplete penetrance of the sex-determining locus has been reported in experimental families of this species (16). Thus, the reliability of the assembly results of the 130-kb male-specific region and one of its adjacent region in Chr19Y was confirmed. Note that we were not able to apply this approach to the 115-kb region due to its highly repetitive nature.

### *k*-mer analysis identifies male-specific sequences in *T. snyderi* and *T. vermicularis*

We used similar strategies to characterize the sex-determining locus of *T. snyderi* on Chr18 and that of *T. vermicularis* on Chr10 (shown in red in Fig. 1A). First, we searched for the male-specific 35-mers in the resequencing data of 24 individuals for each sex of *T. snyderi* (three pools of both females and males, each pool containing eight individuals) and four individuals for each sex of *T. vermicularis* (Table S5), and assembled them into contigs (Table S6). BLAST searches of the contigs against the *T. niphobles* assembly revealed that many of the male-specific contigs in both species showed high similarity to part of the male-specific sequence in *T. niphobles*. For example, the male-specific contigs in both species contain a coding sequence of *GsdfY* (Table S7). This result was unexpected as the chromosomal locations of the sex-determining loci in the three species are distinct from each other, raising the possibility that the male-specific region is shared among the three species, at least in part.

### Construction of the genome assemblies of *T. snyderi* and *T. vermicularis*

We then generated a genome assembly for *T. snyderi* and *T. vermicularis*, including their Y chromosomes. Because individuals with the YY genotype were not available, the genome of an XY male was sequenced for each species. We used Oxford Nanopore long reads and MGI single-tube long-fragment reads (stLFRs) for *T. snyderi*, and PacBio long reads and Illumina short reads for *T. vermicularis* (Table S16, S17, and S18). The male assembly for *T. snyderi* consisted of 1,097 scaffolds with an N50 of 6.0 Mb and a total size of ∼399 Mb, whereas the male assembly for *T. vermicularis* comprised 1,756 contigs with an N50 of 1.57 Mb and a total size of ∼428 Mb (Table 1). A BUSCO search against the 4,584 single-copy orthologs for Actinopterygii suggested that only 1.2% and 2.1% of the core genes were undetected in the genome assemblies of *T. snyderi* and *T. vermicularis*, respectively (Table S12). Approximately 95% of the short reads from male individuals of both *T. snyderi* and *T. vermicularis* mapped uniquely to the respective assembled genomes with a mapping quality above 20.

### The assembled genomes for *T. snyderi* and *T. vermicularis* contain their male-specific regions

To determine the male-specific region in the assembly for *T. snyderi*, we aligned resequencing reads from females and males and compared the relative depth of coverage between them. This analysis revealed a clear difference in coverage between sexes on scaffold_373 (79 kb) (Fig. 3A). A BLAST search revealed conserved synteny between this region and the 130-kb male-specific region harboring *GsdfY* in *T. niphobles* (Fig. 3C, Fig. S10A).

**Fig. 3.**
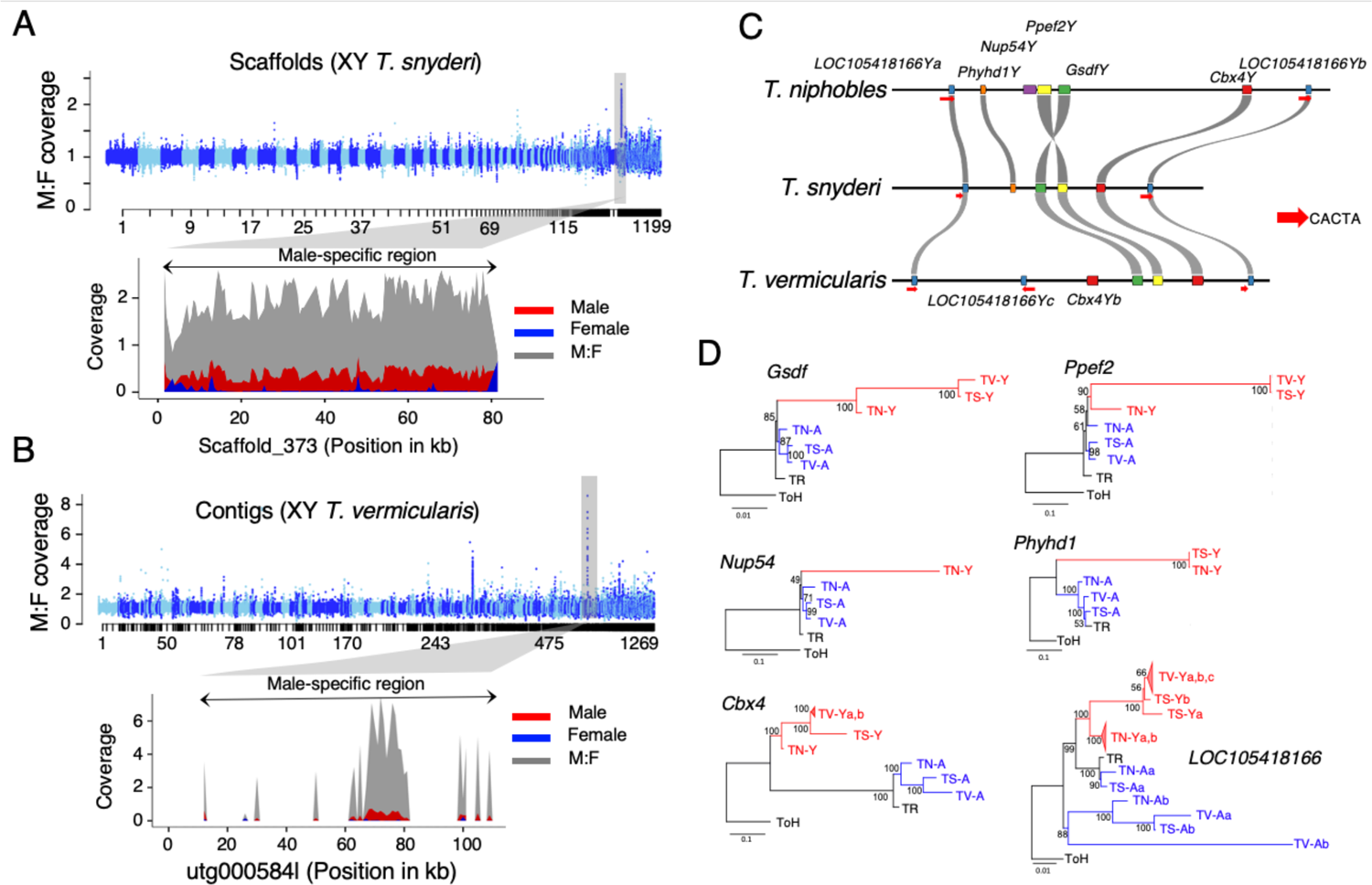
Translocation of the shared male-specific region drove the turnover of sex chromosomes in *Takifugu* species. (A) The male-specific region in *T. snyderi*. We aligned resequencing reads from 24 males and 24 females against the assembled genome of a *T. snyderi* XY male and calculated the relative depth of coverage between sexes ((average male coverage+1) / (average female coverage+1)) at a sliding-window size of 1-kb. In the upper plot showing the genome-wide analysis, scaffolds are on the X-axis. The lower plot shows the male-specific region on scaffold_373. The normalized coverage of males, females, and their ratio are shown in red, blue, and gray, respectively. (B) The male-specific region in *T. vermicularis*. We aligned resequencing reads from 7 males and 12 females against the assembled genome of a *T. vermicularis* XY male. In the upper plot showing the genome-wide analysis, contigs are on the X-axis. The lower plot shows the male-specific region found on contig utg000584l. Note that this contig contains large inverted duplications and a repeat-rich region (Fig. S10B) in which paired-end reads cannot be mapped except for in several segments. The contiguous sequence spanning all the male-specific segments was defined as the male-specific region. (C) Schematic representations of the male-specific regions in *T. niphobles*, *T. snyderi*, and *T. vermicularis*. Each haplotype is annotated with pentagons to depict annotated full-length genes. Arrows indicate CACTA transposable elements containing the catalytic center of a transposase. (D) Maximum-likelihood clustering of the male-specific and autosomal paralogs in *Takifugu niphobles*, *T. snyderi*, and *T. vermicularis*. Red and blue colors represent the male-specific (-Y) and autosomal (-A) paralogs, respectively. TR, TN, TS, and TV represent *T. rubripes*, *T. niphobles*, *T. snyderi*, and *T. vermicularis*, respectively. The autosomal orthologs in *Torquigener hypselogeneion* (ToH) were used as outgroups. Bootstrap values are shown at the nodes of each tree. Source data underlying these panels are provided as a Source Data file.

We used the same approach for *T. vermicularis*, and found coverage differences between sexes on contig utg000584l (97 kb) (Fig. 3B). As in the case of *T. snyderi*, sequence similarity was observed between this contig and the 130-kb male-specific region in *T. niphobles*. In addition, this contig contained large inverted duplications and a repeat-rich region where paired-end reads could not be properly mapped (Fig. S10B).

The association of the male-specific region with phenotypic sex in a wild population of both species (n = 19 and 20 for each sex, respectively, of both *T. snyderi* and *T. vermicularis*) was supported using a diagnostic primer pair that could produce amplicons of different sizes from the male-specific region and its paralogous region (P = 5.7 × 10^-11^ and P = 3.3 × 10^-9^, respectively) (Fig. S9).

### Male-specific regions are largely conserved among the three species

Manual annotation of the male-specific regions in *T. niphobles* (130 kb), *T. snyderi* (79 kb), and *T. vermicularis* (97 kb) revealed similar characteristics of these regions; they harbor *GsdfY* and some other shared genes, and are flanked by two CACTA transposons (Fig. 3C). The sizes of the segments are also similar: 115 kb in *T. niphobles*, 61 kb in *T. snyderi*, and 105 kb in *T. vermicularis*. The gene contents in these regions were found to be similar but distinct. For example, while *GsdfY*, *Ppef2Y*, *Cbx4Y*, *LOC105418166Y*a, and *LOC105418166Yb* were shared among all three species, *Nup54Y* was observed only in *T. niphobles*. Moreover, *T. vermicularis* appeared to have lost *Phyhd1Y* but acquired an additional copy of *Cbx4Y* and *LOC105418166Y* (Fig. 3C, Fig. S10).

To understand the history of the male-specific region that appeared to be conserved but located on different chromosomes in the three species, we constructed phylogenic trees for each gene in the male-specific region and its autosomal paralog(s) across the three species (Fig. 3D). This phylogenetic analysis identified three main branching patterns (Fig. 3D, Fig S11 and S12). The pattern of *Gsdf* indicated that the male-specific region (red in Fig. 3D) and the autosomal paralogs (blue in Fig. 3D) in the three species diverged after their split with the lineage leading to *T. rubripes*, but before the diversification of the three species. The second pattern, including *Ppef2* and *Nup54*, suggests that the topology at the crown age of the genus is ambiguous. The third pattern, including *Cbx4*, *Phyhd1*, and *LOC105418166*, suggested that these paralog pairs diverged before the split of the three species with *T. rubripes*. These results indicate that the male-specific region may have been formed due to capture of *GsdfY* by the pre-existing segment containing other paralogs. Alternatively, the relaxation or intensification of selection for each gene in the male-specific region may underlie the gene topology (39).

Despite the heterogeneous topologies observed in the gene trees, all paralog pairs in the three species were clustered not according to species genealogy, but rather to paralog groups (red in Fig. 3D, Fig. S11-S13), suggesting that the male-specific genes were likely transposed or translocated as a unit into the current genomic positions after the initial formation of the supergene. When focusing on the male-specific paralogs for each species, duplicates of the gene flanking the male-specific region, *LOC105418166Y*s, were clustered according to species group (Fig. 3C). Duplicates of *Cbx4Y* in *T. vermicularis* were also clustered according to species group. Thus, it is likely that the linage-specific duplication of genes occurred in the gene cassette of the male-specific region.

## Discussion

Based on a combination of extensive genetic and genomic data, this study documents evidence that the evolution of the male-specific supergene underlies the replacement of the sex-determining gene in an ancestor of a subset of *Takifugu* fishes, and that subsequent translocation of the supergene drove a rapid sex chromosome turnover without a simultaneous change in the sex-determining gene. Presumably, the accumulation of genes in the male-specific region conferred a selective advantage of this region over the ancestral sex-determining gene, *Amhr2*. However, the insertion of the supergene could have caused sex chromosome differentiation that resulted in a selective disadvantage, namely the accumulation of repeats (toxic Y hypothesis) (35), which may in turn have facilitated translocation of the core part of the supergene at relatively short intervals.

### Formation of the male-specific supergene

Supergenes are sets of tightly linked genes that contribute to a complex phenotype (36–41). Sex-determination regions can be regarded as supergenes because suppressed recombination, if present, causes cosegregation of genes linked to the sex-determining gene (42).

Although direct evidence is lacking in *Takifugu* fish, the genes linked to *GsdfY* in the core male-specific region may play a role in testis differentiation, because either they or their paralogs are expressed in the male gonad during gonadal differentiation (Fig. S6). Moreover, previous studies in other animals have suggested that some genes in the male-specific region of the three *Takifugu* fishes are involved in gonadal differentiation and/or sexual behavior. For example, *Nup54* regulates nucleocytoplasmic transport, whose possible role in gonadal differentiation has been proposed based on its conserved expression in teleost fish (43). In *Drosophila*, this gene is essential in transposon silencing in the ovary (44), and there is evidence suggesting that it plays a role in sexual conflict by influencing female behavior (45). Moreover, this gene is contained in a supergene under sexual antagonistic selection in rainbow trout (46). *Ppef2* is predominantly expressed in photoreceptors and the pineal gland, and it participates in phototransduction in mammals. This gene is also contained in the rainbow trout supergene under sexual antagonistic selection (46). *Cbx4* codes for a member of the Polycomb-group proteins that control cell identity and development (47). Although its role in sexual dimorphism has not been reported, *Cbx2* codes for another member of the group and is required to stabilize the testis pathway in mice (48).

Under some circumstances, males in which the male-specific region was expanded by the addition of these genes were likely to have had a selective advantage over males that retained the ancestral sex-determining gene. Interestingly, an observation that appears to support this hypothesis is that the majority (75.6% of 131 fish) of the wild F1 hybrids between *T. snyderi* and *T. stictonotus* were derived from *T. snyderi* males (49). Since *T. snyderi* males carry the supergene containing *GsdfY* while *T. stictonotus* males carry the Y allele (G allele at the SNP site) of *Amhr2* (see Fig. 1A and 1B), it is tempting to speculate that the difference in sex-determining loci underlies the asymmetric reproductive success.

### The evolutionary development of the core male-specific region

Although supergenes contribute to many fascinating phenotypes (11–14), their evolutionary development is often only incompletely understood. In particular, the analysis of sex-determining regions has been difficult due to the extensive accumulation of repetitive sequences (see some exceptions in (24–28)). Our study provides a unique opportunity to infer the process involved in assembling the male-specific supergene and its subsequent modifications during repeated translocation.

It is likely that the core male-specific region evolved in the common ancestor of the three species after diverging from the lineages leading to *T. rubripes* and *T. obscurus* through the combination of at least two processes. One includes segmental duplication encompassing *Nup54, Ppef2*, and *Gsdf* on Chr6 and the translocation of this region to the future sex chromosome (Fig. S13 and S14), given that the genomic arrangement of the three genes adjacent to each other is a conserved feature among teleost fish (43). The other process is the gathering of other unlinked genes (*Phyhd1*, *Cbx4*, and *LOC105418166*) through independent duplications and translocations (Fig. 3C and Fig. S14). It is possible that the latter process partly predates the former, as the phylogenetic analysis in this study implied that the emergence of *Phyhd1Y* and *LOC105418166Y* occurred prior to that of *GsdfY*. The original segment, if this is the case, was probably not male specific until it captured *GsdfY*.

Based on the core structure formation, it is parsimonious to consider that *Nup54Y* was lost before the divergence of *T. snyderi* and *T. vermicularis*, while the gene content was retained in the lineage leading to *T. niphobles* (Fig. 1A and Fig. S14). Then, in the *T. vermicularis* lineage, while *Phyhd1Y* was lost, the segmental duplication of part of the male-specific region resulted in the emergence of *LOC105418166Yc* and *Cbx4Yb* (Fig. 3C, Fig. S10 and S14). The variations in gene content in the shared male-specific region suggest that adaptive replacements of genes may have taken place in these species and could still be ongoing. The polyphyletic relationship of the male-specific supergene in the three species (Fig. 1A) could be due to incomplete lineage sorting in the rapid speciation of *Takifugu*.

### The jumping sex-determining locus

In vertebrates, transposition or translocation of a pre-existing sex-determining locus has only been reported in salmonids (50). Comparative analyses of the sex-determining loci in the three salmonid species (51) suggested that an ∼4.1-kb sequence containing the sex-determining gene *sdY* is conserved, along with transposable elements. Although an RNA-mediated mechanism has been proposed for the transposition of the *sdY* locus in salmonids, different mechanisms likely played a role in the movement of the male-specific region in *Takifugu*, as the size of the supergene spans ∼100-kb. A notable feature of the gene arrangement of the conserved male-specific region is that the region is flanked by two duplicated regions harboring CACTA transposable element and *LOC105418166Y* (Fig. 3C). Therefore, the flanking transposons are candidate drivers for relocating the male-specific region. However, we were unable to find the terminal inverted repeats (TIRs) that are expected to reside at the ends of transposons if a cut-and-paste process is involved in transposition. Although it is possible that the TIRs have degenerated, the same orientation of the two transposons is incompatible with the standard model of cut-and-paste transposition (32). Based on the orientation of the two flanking regions, we prefer the hypothesis that non-allelic homologous recombination played a role in the translocation. This process is thought to contribute to duplication and translocation events in animals and plants (52–54), and suggests that recombination between two directly oriented homologous sequences (typically, repeats or transposable elements) can lead to deletion and insertion of the genomic region flanked by these sequences (55). As a result, duplicated and translocated fragments are frequently bordered by transposable elements (56). Of note, this model is only speculative in *Takifugu*, and the mechanism underlying the translocation of supergene is still undetermined.

### Roles of translocations in accumulation and purging of deleterious sequences

It is hypothesized that sex chromosomes start to differentiate when recombination around the sex-determining region stops. While the accumulation of deleterious mutations on the differentiating sex chromosomes will eventually reduce fitness (6, 35, 57, 58), the “jumping” ability of the sex-determining locus is a way to purge the deleterious mutations (50). Consistent with these hypotheses, our analysis of *k*-mers and read coverage suggested that half of the male-specific region (115 kb), which contains many large repetitive sequences in *T. niphobles*, appeared to be purged from *T. snyderi* and *T. vermicularis* after the translocations. The 115-kb region may have arisen due to the reduced recombination triggered by the insertion of the 130-kb region containing *GsdfY* in an ancestor of the three species. However, it is possible that the 115-kb region has recently emerged in *T. niphobles* in a species-specific manner. The distinction of the two scenarios was challenging due to the presence of multiple duplicated loci corresponding to genes in the *T. niphobles* 115-kb region in the genomes of *Takifugu* fishes, which made it difficult to distinguish orthologs and paralogs. To fully evaluate the consequence of repeated translocation of the sex-determining supergene in the three species, more contiguous sequences of the genomes for *T. snyderi* and *T. vermicularis* are needed.

### Remarks

This study uncovered a process involved in the replacement of both sex chromosomes and sex-determining genes in closely related species, and highlighted the importance of repeated translocation of the sex-determining region. Moreover, the construction of contigs for the jumping sex-determining supergene revealed that it has a unique structure flanked by two putative transposable elements. Outside of vertebrates, the translocation or transposition of a pre-existing sex-determining gene was recently characterized in insects (59) and plants (60, 61). The repeated translocation of a sex-determining locus may be more prevalent in vertebrates than previously thought, and may have caused rapid turnover of sex chromosomes.

## Materials and methods

### Conservation of the sex-determining SNP in *Amhr2*

The genotype at SNP7271 on *Amhr2* is associated with phenotypic sex in fugu (15, 16). We previously examined whether the association is conserved in *T. pardalis*, *T. poecilonotus*, and *T. niphobles*, and found that while it was identified in *T. pardalis* and *T. poecilonotus*, the locus was fixed with the female allele (C allele) in *T. niphobles* (15, 16). In this study, we extended the analysis to eight *Takifugu* species, namely *T. chrysops*, *T. stictonotus*, *T. obscurus*, *T. ocellatus*, *T. xanthopterus*, *T. porphyreus*, *T. snyderi*, *and T. vermicularis*, and genotyped a total of 342 wild-caught and 15 aquacultured fish by a direct-sequencing approach. The genomic region containing exon9 of *Amhr2* and its adjacent regions was PCR amplified and directly sequenced as previously reported (16). SNP loci were identified and genotyped based on the sequence data. Additional genotyping of SNP7271 of *T. snyderi* (*n* = 100) and *T. vermicularis* (*n* = 100) was performed by high-resolution melting analysis using a Lightscanner (Idaho Technology, Salt Lake City, UT, USA) following the method of (22). Information on the sampling locations, sex, and association analysis (Fisher’s exact test) results of the fish of each species is provided in Table S1. No specific permission was required for this research as the samples were obtained through official fishermen’s unions. This study did not involve endangered or protected species.

### Linkage mapping

#### Experimental families of *T. snyderi*

Three families (Families A, B, and C) of *T. snyderi* were produced between three wild males and a single wild female caught at Suruga Bay, Shizuoka Prefecture, Japan (Table S2). Procedures for artificial fertilization and rearing conditions were defined as previously reported for *T. niphobles* (62). Fish at 122 and 174 days post fertilization (dpf) were combined and used for the following analyses. Phenotypic sex was visually determined under a microscope.

#### Genotyping of SNP markers in *T. snyderi*

GRAS-Di (Genotyping by Random Amplicon Sequencing–Direct) technology (63, 64) was used to obtain the genome-wide genotype data from 98 fish in Family A. Genomic DNA was extracted from the caudal fin using the Gentra Puregene tissue kit (Qiagen, Hilden, Germany). Library construction and sequencing were conducted by Eurofin Genomics as described in (64). The sequencing was performed on an Illumina HiSeq 2500 platform (PE 100 bp). Reads were trimmed using Trimmomatic (v. 0.36) (65) as described in (64), and mapped onto a reference genome sequence of *T. rubripes* (fugu), FUGU5 (23) (GenBank: GCA_000180615.2) using BWA-MEM (66) with the default parameters. SNP calling was performed using GATK HaplotypeCaller (v. 3.8) (67) with the default setting. VCFtools (v. 0.1.14) (68) was used to extract SNPs meeting the following criteria: minor allele frequency between 0.1 and 0.4, genotyped for ≥ 90% of individuals, allele count of two, and minimum depth of 10. The SNP loci heterozygous in both parents were removed. The sequence data were registered in the DDBJ SRA (Accession No. DRA012888) The genotype information is provided in Supplementary material Gras-Di Mapping for Family A in “genotype data.xlsx.”

### Linkage map construction and mapping of the sex-determining locus in *T. snyderi*

The male and female linkage maps were constructed based on the paternally and maternally inherited alleles at marker loci, respectively, using the R/qtl package (v. 1.41) (69). Linkage groups were inferred with the *formLinkageGroups* function (max.rf = 0.45, min.lod = 6.0), and the markers were ordered by the *orderMarkers* function. Linkage groups with fewer than five loci were excluded from the subsequent analysis. The sex-determining locus was analyzed by the interval mapping implemented in R/qtl as described in (16). The genome-wide significance level was determined with a permutation test with 10,000 permutations (70). The chromosome and linkage group identities of *T. snyderi* were assigned based on synteny with those of *T. rubripes* established in (23).

#### Fine mapping of the sex-determining locus using microsatellite markers in *T. snyderi*

Although interval mapping is a powerful tool to identify loci underlying differences in phenotypes, it is possible that the genotype–phenotype association observed at the distal end of Chr18 is family specific. To test this possibility, we used 83, 25, and 62 fish from Families A, B, and C, respectively, and genotyped markers f1618 and f1659, which were previously mapped near the distal end of Chr18 in the fugu genome. The association between gonadal sex and genotype was assessed by Fisher’s exact test (Table S3). To increase the resolution, we genotyped two marker loci (f1335 and f1433) on Chr18 for 765 fish from the three families (342, 158, and 265 fish in Families A, B, and C, respectively). After identifying 30 recombinants, 26 microsatellite markers flanked by the two markers on Chr18 were genotyped for these individuals (Fig. S1). Genotyping procedures are described in (23). Information on the markers is provided in Table S19.

#### Experimental families of *T. vermicularis*

For *T. vermicularis*, two pairs of wild-caught parents from Ariake Sea (Nagasaki Prefecture, Japan) were mated independently to produce two families, Families D and E (Table S2). Fish production and phenotyping were performed as in the case of *T. snyderi*.

### Linkage map construction and mapping of the sex-determining locus in *T. vermicularis*

We first conducted a genome-wide linkage analysis of the sex-determining locus using 62 fish at 127 dpf in Family D with 138 genome-wide microsatellite markers (Table S2). The linkage between phenotype and genotype observed at the distal end of Chr10 in Family D was confirmed using 164 fish in Family E sampled at 131–161 dpf by genotyping 35 markers anchored on Chr10 in *T. rubripes*. The genotype information is provided in Table S20 and Mapping for Family D in “genotype data.xlsx.”

### The phylogenetic framework of *Takifugu*

To generate a species phylogenetic tree, whole-genome resequencing data were obtained from 12 *Takifugu* species and one outgroup (one individual per species): *Takifugu rubripes*, *T. snyderi, T. vermicularis, T. niphobles, T. pardalis, T. poecilonotus, T. chrysops, T. stictonotus, T. obscurus, T. ocellatus, T. xanthopterus, T. porphyreus,* and *Torquigener hypselogeneion* (outgroup). A detailed description of the method used to generate the species tree is provided in the supplementary methods.

### *k*-mer analysis to identify male-specific sequences

A detailed description of k-mer analyses of *T. niphobles*, *T. snyderi*, and *T. vermicularis* is provided in the supplementary methods.

### Genome assembly in a *T. niphobles* YY male Long- and short-read sequencing

We sequenced the genome of a YY individual on a PacBio Sequel platform. The production of the YY fish was reported previously (16). In brief, we first identified an XY sex-reversed female using sex-linked microsatellite markers (f2003 and f2006) to cross with an XY male. The offspring consisted of individuals carrying XX, XY, or YY sex chromosome sets. Then the YY fish was selected based on the sex-linked markers f2003 and f2006. Genomic DNA of the YY fish was isolated from the caudal fin by the conventional phenol/chloroform/isoamyl-alcohol method. Size selection, library preparation, and sequencing were conducted at the National Institute of Genetics. Approximately 56.6-Gb sequences with total reads of 5,445,262 (N50 = 17 kb) were derived from eight PacBio SMRT cells (Table S8). Genomic DNA from the YY fish was also used for whole-genome Illumina resequencing. A TruSeq DNA PCR-Free library was prepared and sequenced (2 × 245 bp) on the HiSeq 2500 platform at the National Institute of Genetics. The data have been registered in the DDBJ SRA (Accession No. DRA012992).

### Hybrid genome assembly and Hi-C scaffolding

Illumina reads were trimmed with Trimmomatic (v. 0.36) (65) using the following parameters: ILLUMINACLIP TruSeq3-PE-2.fa:2:30:10, LEADING:10, TRAILING:10, SLIDING WINDOW:30:20, AVGQUAL:20, and MINLEN:240, and the resulting reads were assembled with SparseAssembler (v. 20160205) (71) into contigs with k-mer size = 51 and skip length = 15. The contiguity of the short-read assembly was improved by anchoring the contigs to the PacBio long reads using DBG2OLC (v. 20180222) (72) with *k*-mer size = 17, minimum overlap = 30, and AdaptiveTh = 0.01. After removing chimeric reads with the option of *ChimeraTh* = 1, we obtained an initial hybrid assembly. Because raw PacBio long reads are prone to contain errors, we corrected potential sequencing errors in the initial assembly by realigning the long reads and the Illumina contigs using Sparc (v. 20160205) (73). We also aligned the long reads to the assembly with pbalign (v. 0.4.1) (https://github.com/PacificBiosciences/pbalign/) using the BLASR (v. 5.3.1) (74) algorithm and polished it with Arrow (v. 2.3.3) with the –minCoverage 12 option (75). For error correction at the base level, Illumina reads were aligned to the assembly using BWA-MEM (66), and the assembly was polished three times with Pilon (v. 1.22) (76). For long-read-only assembly, the Minimap2/Miniasm pipeline (77, 78) was used. In brief, a Pairwise mApping Format (PAF) file was generated using Minimap2 with the -x ava-pb option. The PAF file was converted into an assembly graph as a Graphical Fragment Assembly (GFA) format file using Miniasm. Then, a *de novo* assembly sequence (unitig) was extracted from the GFA file. The long reads were mapped to the unitig sequences using Minimap2, and three rounds of consensus corrections were performed using Racon (v. 1.3.0) (79). The resultant assembly was polished three times with Pilon as described above. Finally, the hybrid assembly and long-read-only assembly were merged using Quickmerge (v. 0.2) (80), employing the latter as the query assembly. The merged assembly was then merged to the long-read-only assembly to produce a more contiguous assembly. Furthermore, Hi-C data from an XY male individual were used to scaffold this merged genome. Details of tissue collection, Hi-C sequencing, and Hi-C scaffolding are provided in the Supplementary methods. An overview of the genome assembly is shown in Fig. S2.

### Assembly quality assessment

The quality of the assemblies was assessed by gVolante (v. 1.2.1) (81) with BUSCO (v. 3.0.2) (82) and the ortholog gene set of Actinopterygii. In addition, the quality of the Hi-C–based proximity-guided assembly was evaluated by mapping the Illumina short reads (sample IDs: 19m, 12m, 10m, 11m, and 0525m in Table S5) to the assembled genome using BWA-MEM with default parameters. A comparison of this genome assembly with that of *T. rubripes* (FUGU5) was also conducted (see details in the supplementary methods).

### Gene annotation

Gene annotation was conducted using MAKER (v. 3.01.02) (83), which utilizes both evidence-based methods and ab initio predictions. For the evidence-based annotation, the RNA-seq data of XY *T. niphobles* and the annotated protein sequences from FUGU5 were used with default parameters. RNA-seq data of XY *T. niphobles* were generated on an Illumina HiSeq 2000 platform at Novogene (Tokyo, Japan). Sample collection, RNA extraction, library preparation, and sequencing strategies are described in the supplementary methods. The ab initio annotation was performed using SNAP (84), trained on the gene model from the evidence-based annotation, and AUGUSTUS (85), trained on the Actinopterygii gene model (82) from BUSCO (v. 3.0.2). The curated libraries of repeats for *T. niphobles* were also used for the annotation (see details of the repeat annotation in the supplementary methods).

#### Samples and additional resequencing of *T. niphobles*

For a comparison of the relative depth of read coverage between males and females, we sequenced an additional two females and four males of *T. niphobles* from Lake Hamana. Phenotypic sexing and genomic DNA extraction were carried out as described above. Library preparation and sequencing on HiSeq 2000 platform were performed by BGI Corporation (Kobe, Japan). Information about the sequencing statistics is presented in Table S5. These reads were trimmed by removing adapters and low-quality reads as described above, with either the MINLEN:80 (m8, m, m30, and m31) or MINLEN:140 (35fe and 11fe) option. The data were registered in the DDBJ SRA (Accession No. DRA012877).

### Comparison of the relative depth of read coverage between males and females

To determine the male-specific region in the genome assembly of the *T. niphobles* YY male, we mapped the resequencing data of 10 males and 8 females (Table S5) onto the *T. niphobles* reference sequence using BWA-MEM (66) with default settings, and compared the relative depth of coverage between sexes. To count the depth, we used Mosdepth (v. 0.3.0) (86) with a 1-kb window size and excluded regions with depth greater than 70x. Mean coverage values were calculated separately for females and males, after normalization of each sample with the genome coverage. The male-to-female coverage ratio was calculated as (average male coverage+1) / (average female coverage+1) and plotted along 22 chromosomes using the R/qqman package (87).

### Characterization of the male-specific region

To identify duplicate and satellite sequences within the male-specific region, the sequences of the male-specific region were aligned against themselves using BLASTn with the “word_size 50” option. Aligned blocks longer than 50 bp were visualized by ggplot2 (v. 3.3.5) (88). To test if the male-specific region comprised segments that were duplicated and translocated from other regions of the genome, we first performed a BLAST search of the male-specific region against the reference genome of *T. niphobles*, and then aligned the male-specific region to the genome assembly of *T. niphobles* using the nucmer algorithm with option -l 50 -c 100 implemented in MUMmer (v. 4.0) (89). Synteny blocks longer than 1-kb with 95% sequence identity were visualized by Circos Plot using circos (v. 0.69-7) (90). We verified the contiguity of the assembled sequence by tiling PCR. Moreover, the presence of the male-specific region in the wild population of *T. niphobles* was confirmed by a diagnostic primer. A detailed description of these strategies is presented in the supplementary methods.

### Genome assembly in a *T. snyderi* XY male Long- and linked-read sequencing

The whole-genome sequence of an XY individual from Lake Hamana was obtained using PromethION (Nanopore Technologies) and MGI single-tube long-fragment reads (stLFRs, MGISEQ-2000RS) (Table S16 and S17). Genomic DNA was isolated as described above and the XY genotype was confirmed with diagnostic primers prior to sequencing, according to the method described in the supplementary methods for confirmation of the male-specific region in the wild population. To enhance recovery of DNA fragments longer than 10-kb, the Short Read Eliminator XS kit (Circulomics, Baltimore, MD, USA) was used. Library preparation and sequencing for both technologies were performed by GeneBay (Yokohama, Japan). As for PromethION sequencing, the raw signal intensity data were used for base calling using Guppy (v. 3.0.3). In total, ∼9.09-Gb data of 890,724 (N50 = 25 kb) reads were generated. Low-quality reads (with a mean quality score less than 7) and the adapters were removed using fastp (91) and Porechop (v. 0.2.3) (https://github.com/rrwick/Porechop), respectively. The retained reads were corrected in Canu (v. 1.7.1) (92) with the following settings: minimum read length = 5,000; minimum overlap length = 1,200; and corMinCoverage = 0. This resulted in a total of 7.05-Gb corrected reads. As for stLFR sequencing, the MGIEasy stLFR Library Prep Kit was used to prepare co-barcoding DNA libraries (93). The libraries were sequenced using the MGISEQ-2000RS. In total, ∼38-Gb (without barcodes 32.17 Gb) of paired-end sequences was generated. The stLFR reads were filtered using SOAPfilter (v. 2.2) (94) with a Phred score ≤10 to remove adapter sequences and low-quality reads containing more than 60% bases.

### Hybrid genome assembly and identification of the male-specific region

We generated an initial genome assembly of *T. snyderi* using Canu (v. 1.7.1) (92) with Nanopore long reads. For consensus correction, Nanopore long reads were mapped to the unitig sequences by Minimap2 (78), and three rounds of corrections were performed using Racon (79) followed by one round of correction using medaka (v. 1.2.1) (https://nanoporetech.github.io/medaka/). To improve the Nanopore assembly with stLFR data, we first transformed the stLFR data to pseudo-10X Genomics reads using the stlfr2supernova pipeline (https://github.com/BGIQingdao/stlfr2supernova_pipeline). Then the Nanopore assembly was scaffolded with the transformed stLFR reads using ARKS (v. 1.0.6) with LINKS (v. 1.8.7) (95, 96), using default parameters. The resultant assembly was polished twice using POLCA (97) with the transformed stLFR data. The gene functional annotation of the assembly and its completeness assessment were conducted as described for *T. niphobles*. For the identification of the male-specific region, the relative depth of coverage between males and females was analyzed using 24 individuals for each sex (three pools each of females and males, with each pool containing eight individuals) as described for *T. niphobles*. The characterization of the male-specific region was conducted as described for *T. niphobles*.

### Genome assembly in a *T. vermicularis* XY male

#### Long-read sequencing for genome assembly and short-read sequencing

We sequenced the genome of an XY individual from Ariake Sea using PacBio long reads and Illumina paired-end short reads (Sample ID: S9, Table S5). The genomic DNA was extracted as described for *T. niphobles* (Table S18). The XY genotype was confirmed with diagnostic primers following the aforementioned method for *T. niphobles* (Table S15). DNA fragments shorter than 10-kb were removed using the Short Read Eliminator XS kit. Library preparation and sequencing for PacBio Sequel II were performed at Macrogen Corporation (Kyoto, Japan). In total, ∼128.8-Gb data composed of 11,257,208 subreads (N50 = 15 kb) were generated from a single PacBio SMRT cell. The PacBio subreads were corrected by Canu (v. 1.7.1) with the following settings: minimum read length = 7,000; minimum overlap length = 1,200; and coreOutcoverage = 200.

This resulted in ∼67.17-Gb of corrected reads. For Illumina paired-end short reads, library preparation and sequencing were performed using the NovaSeq 6000 platform of GeneBay. We also sequenced additional eight female and one male *T. vermicularis* from Ariake Sea using the NovaSeq 6000 platform of GeneBay following the aforementioned protocol. Information about the resequencing statistics is presented in Table S5. The short reads were trimmed by removing adapters and low-quality reads as described above, except that the option MINLEN:140 was used.

### Genome assembly and identification of the male-specific region

We generated an initial genome assembly of *T. vermicularis* from the error-corrected PacBio reads using Minimap2 and Miniasm (77, 78) as described for *T. niphobles*. Initial error corrections of the resultant assembly were performed as described for *T. niphobles*. Furthermore, the assembly was polished twice with POLCA (97) using the filtered Illumina reads obtained from a single individual (Sample ID: S9, Table S5). Sequence annotation of the assembly and assessment of its completeness were conducted as described for *T. niphobles*. For identification of the male-specific region, the relative depth of coverage between males and females was analyzed using 7 males and 12 females as described for *T. niphobles*. Characterization of the male-specific region was conducted as described for *T. niphobles*.

### Phylogenetic analysis of male-specific genes and their autosomal paralogs

To determine the phylogenetic relationships of the male-specific genes and their autosomal paralog(s), we used RAxML (v. 0.8) (see details in the supplementary methods).

## Acknowledgments

This work was in part supported by MEXT KAKENHI (No. 221S0002, 24380102, 15H04542, 18H02277, 16K14966, 18K05815, and 17H06425). This work was also supported by the Cooperative Research Grant of the Genome Research for BioResource, NODAI Genome Research Center, Tokyo University of Agriculture and JST-Mirai Program (JPMJMI18CH and JPMJMI21C1).

---

**Fig. S1.**
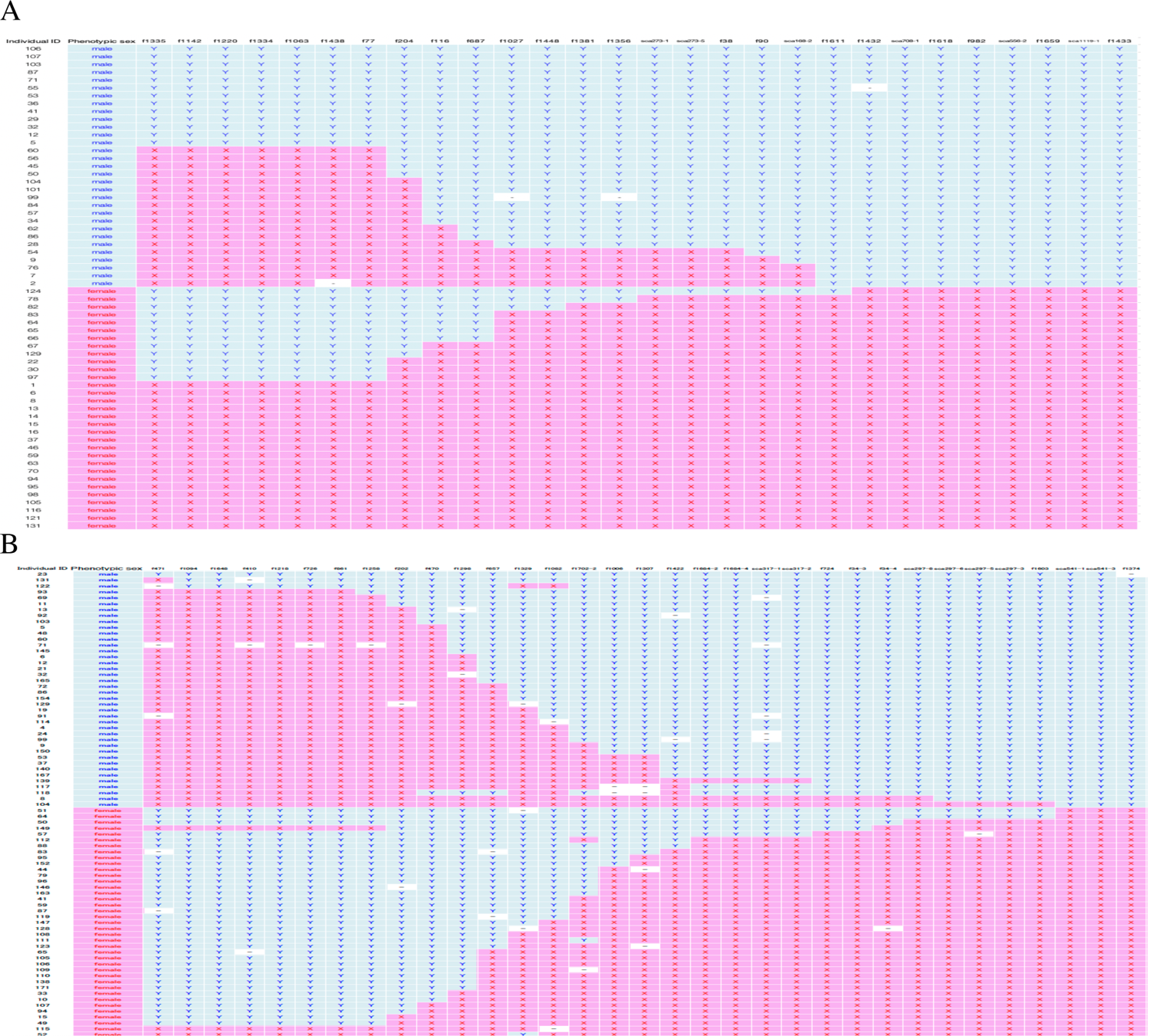
The search for individuals with recombination near the sex-determining locus in *T. snyderi*. (A) and *T. vermicularis* (B). (A) To fine map the sex-determining locus in *T. snyderi*, we first used two markers, f1335 and f1433, that were previously mapped on Chr18 in *T. rubripes* (fugu). We genotyped the two markers in 765 fish from three families (342, 158, and 265 fish in Families A, B, and C, respectively), and identified 30 recombinants. We then genotyped 26 microsatellite markers flanked by the two markers on Chr18 in those individuals. The first row of the table contains the marker names. The rows below show the data of recombinant individuals identified by the screening. “X” and “Y” indicate female- and male-associated alleles, respectively, inherited from the father. Empty blocks indicate that genotypes are not assigned. The results suggest tight linkage of the sex-determining locus with markers near the distal end of Chr18, as well as linkage between the marker loci. Thus, the synteny of this region is conserved between *T. snyderi* and *T. rubripes* (fugu), except for the sex-determining locus. (B) For *T. vermicularis*, 35 markers, most of which have been previously mapped on Chr10 in *T. rubripes*, were used to screen 164 fish in Family E. We found 30 fish with recombination between the most distant markers, f471 and f1374. This analysis suggests both tight linkage of the sex-determining locus with the distal end of Chr10 and conserved synteny of this region between *T. vermicularis* and *T. rubripes*, except for the sex-determining locus.

**Fig. S2.**
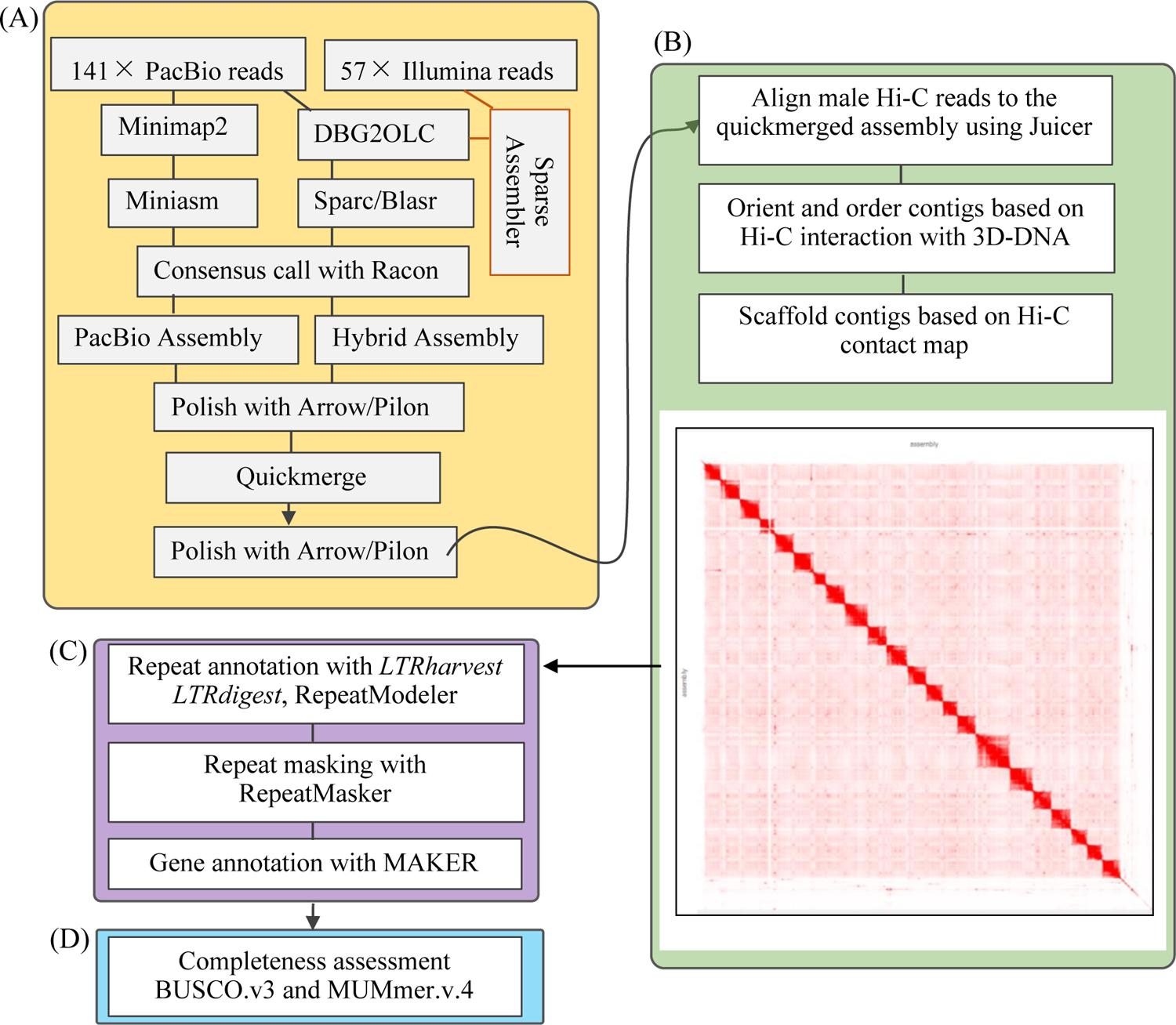
Schematic overview of the genome assembly process in *T. niphobles* with the YY genotype. (A) Initial genome contigs were generated by two strategies: (1) PacBio long-read-only assembly using Minimap2 and Miniasm, and (2) hybrid assembly using DBG2OLC and SparseAssembler by the combination of PacBio long reads and Illumina short reads. The two assemblies were merged using Quickmerge. (B) The Hi-C reads were aligned to the merged assembly using Juicer. Hi-C scaffolding was performed with 3D-DNA using the Juicer output. (C) Gene annotation was conducted using MAKER with evidence from RNA-seq transcriptomes data, predicted protein sequences from FUGU5, and ab initio gene predictions from SNAP and AUGUSTUS, followed by repeat annotation using *LTRharvest*, *LTRdigest*, and RepeatModeler. Repeat-rich regions were identified by RepeatMasker. (D) Genome completeness was assessed by BUSCO (v. 3.0.2).

**Fig. S3A.**
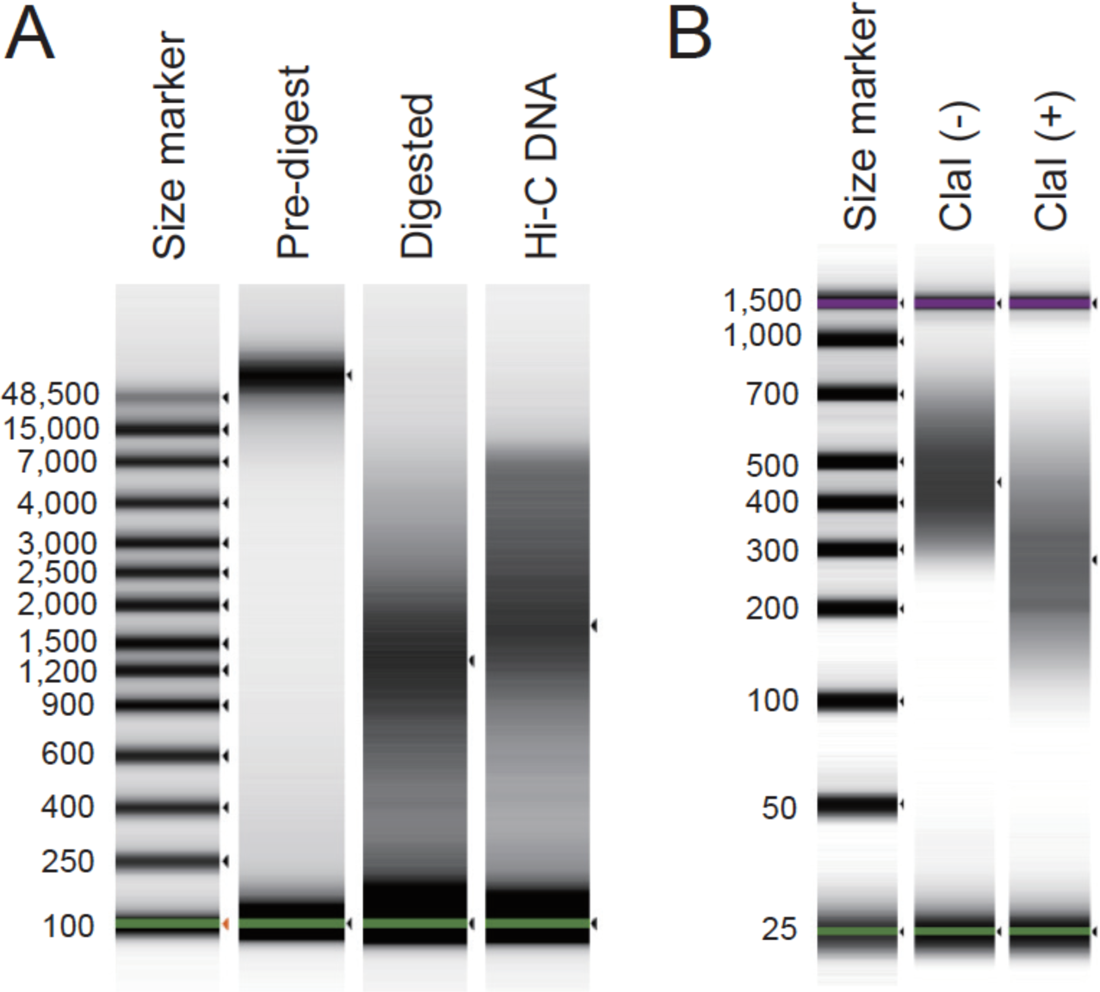
Quality control of Hi-C DNA and the Hi-C library. (A) Size-shift analysis of pre-digested, digested, and ligated DNA (Hi-C DNA) with an Agilent TapeStation using the Genomic DNA ScreenTape assay. (B) Size-shift analysis of the Hi-C library, with or without ClaI restriction enzyme digestion, with an Agilent TapeStation using the High-Sensitivity D1000 ScreenTape assay.

**Fig. S3B.**
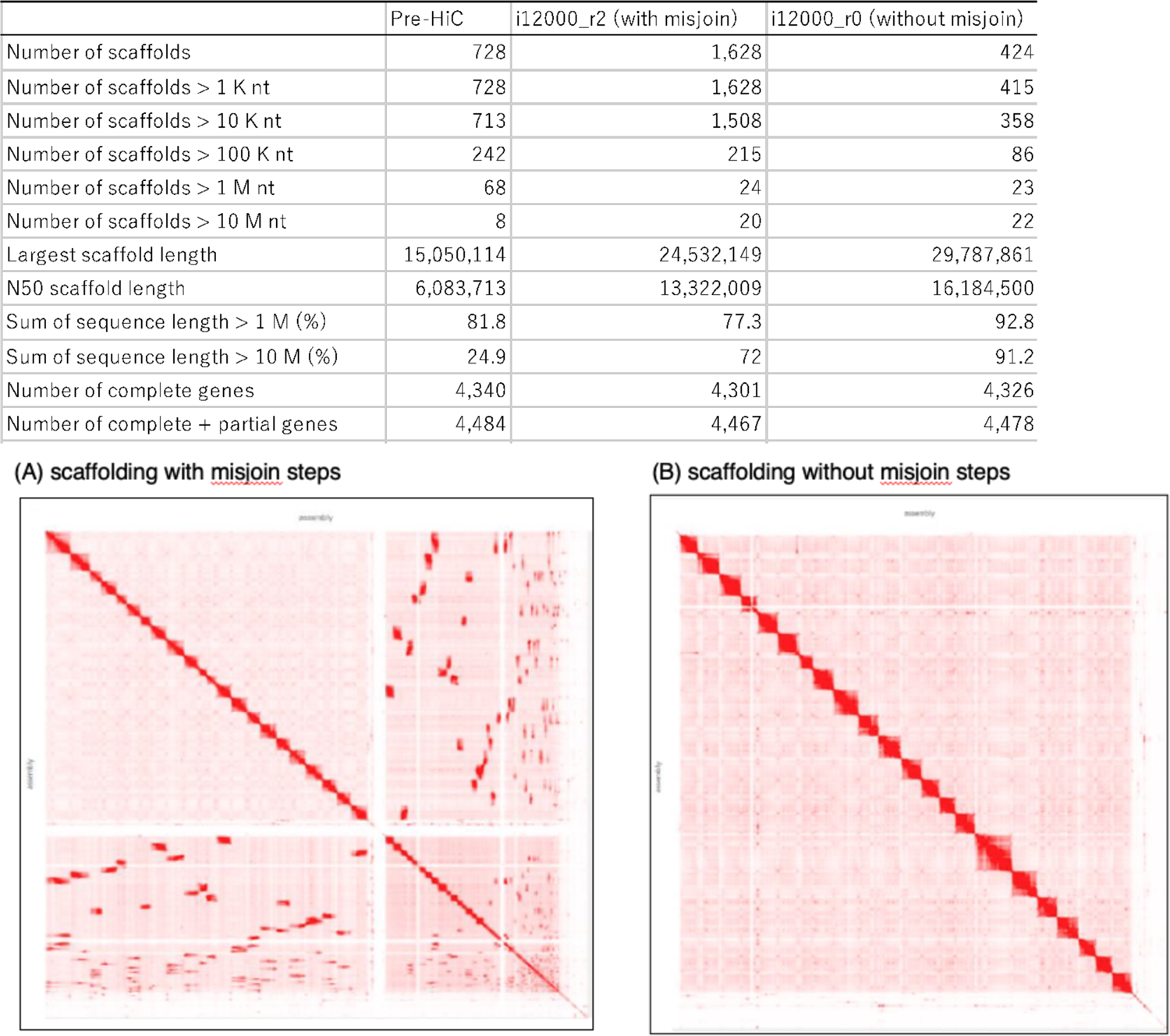
Hi-C scaffolding increased N50 lengths and gene space completeness scores. The table shows a summary of the assemblies before scaffolding, after scaffolding with misjoin steps, and after scaffolding without misjoin steps. Disabling misjoin correction resulted in acceptable chromosome-scale genome sequences, which was validated by the formation of larger and fewer blocks in the Hi-C contact maps (scaffolding with (A) and without (B) misjoin steps), and in an increase in gene space completeness scores (table). In the final assembly, scaffolds of 10 Mb or longer comprise > 90% of the whole genome length, and the number of scaffolds matches the number of chromosomes (n = 22) in *T. niphobles* (1).

**Fig. S4.**
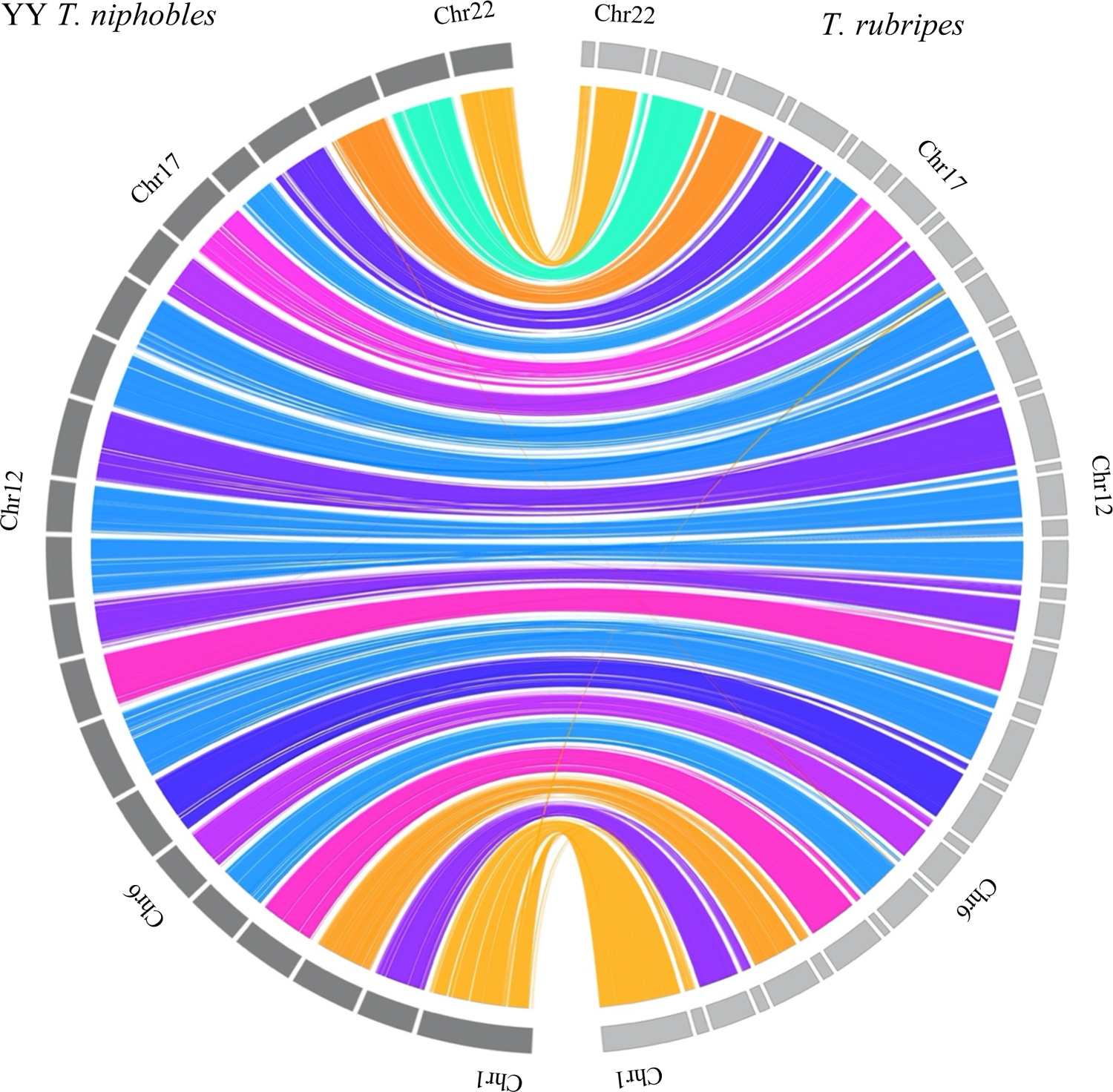
Conserved synteny between the assembled genome of *T. niphobles* with the YY genotype and a reference genome sequence of fugu, *T. rubripes* (FUGU5). Based on conserved synteny, chromosome identities for the assembled genome of *T. niphobles* were assigned according to those of *T. rubripes*.

**Fig. S5.**
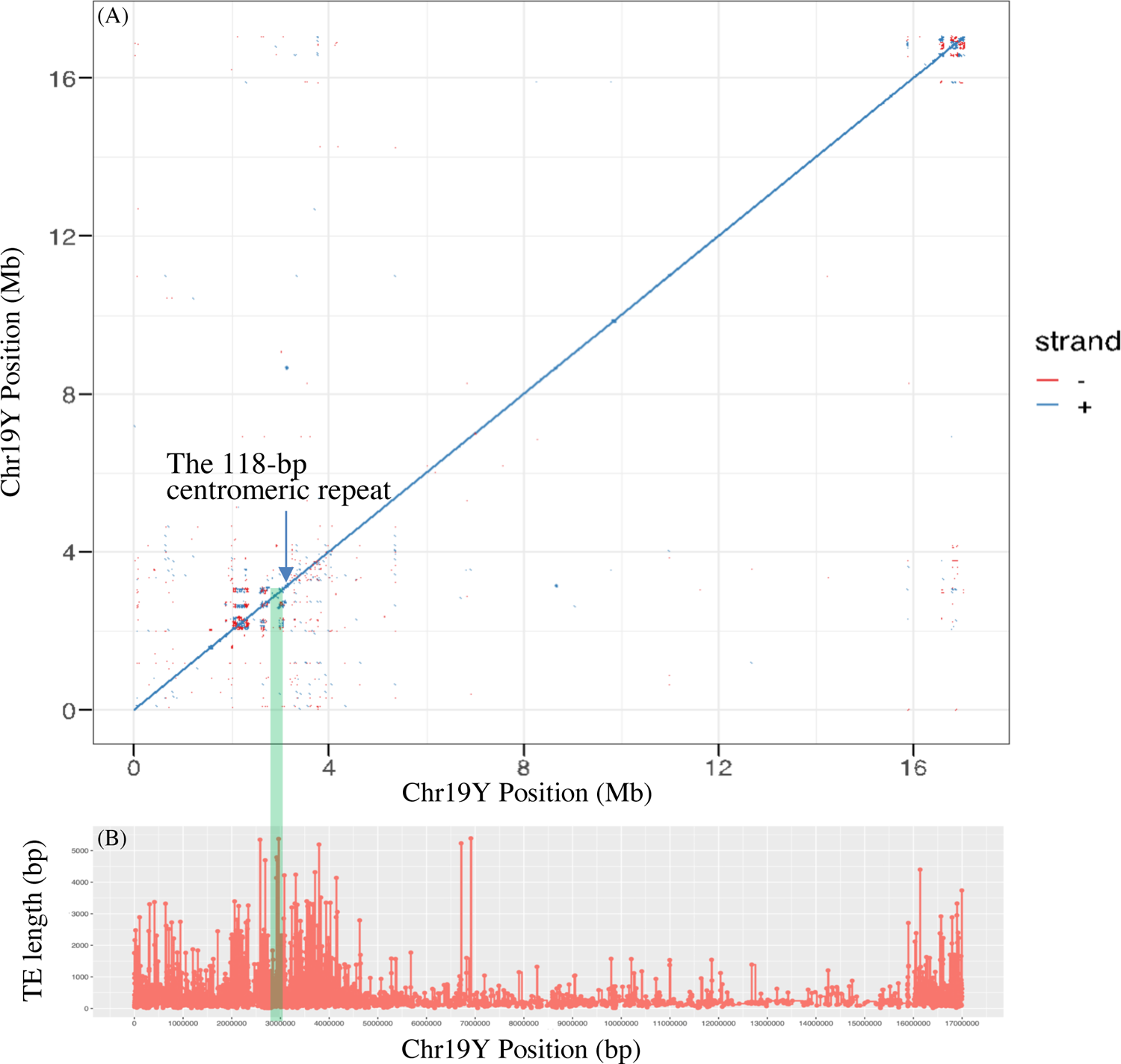
(A) Dot plot of the self-comparison of Chr19Y. The presence of direct (blue) and inverted (red) repeats are visualized as the accumulation of dots off the central diagonal line. The blue arrow indicates the accumulation of a centromeric repeat. The green shadow represents the male-specific region (245 kb). Genomic position is scaled in Mb. (B) The TE length distribution across Chr19Y. Y-axis is scaled in bp.

**Fig. S6.**
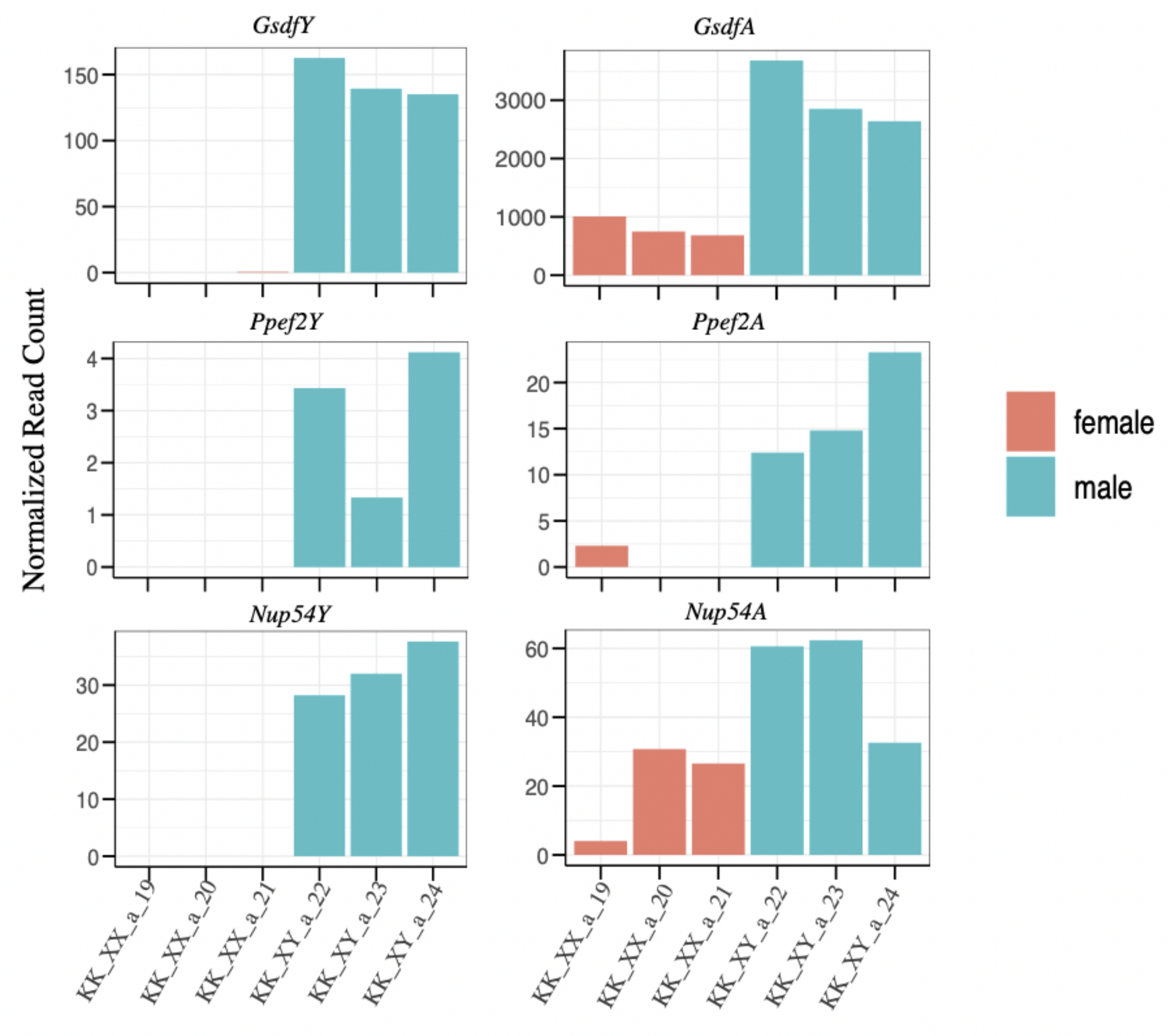
Expression of the genes in the male-specific region and their autosomal paralogs in the developing gonads. We first identified diagnostic single-nucleotide sites on the coding region of *Gsdf*, *Ppef2*, and *Nup54* that could distinguish the male-specific and autosomal paralogs. We then aligned RNA-seq reads from developing gonads (90 dpf) on the reference genome of *T. niphobles* and counted the number of reads mapped on the diagnostic sites. The paralog-specific read counts across libraries were normalized for the library size (total number of paired mapped reads) and expressed per 23 M reads. The candidate sex-determining gene *GsdfY* was expressed in the developing gonads in males only. In addition, autosomal *GsdfA* was more highly expressed in males than in females (three libraries each of females and males).

**Fig. S7.**
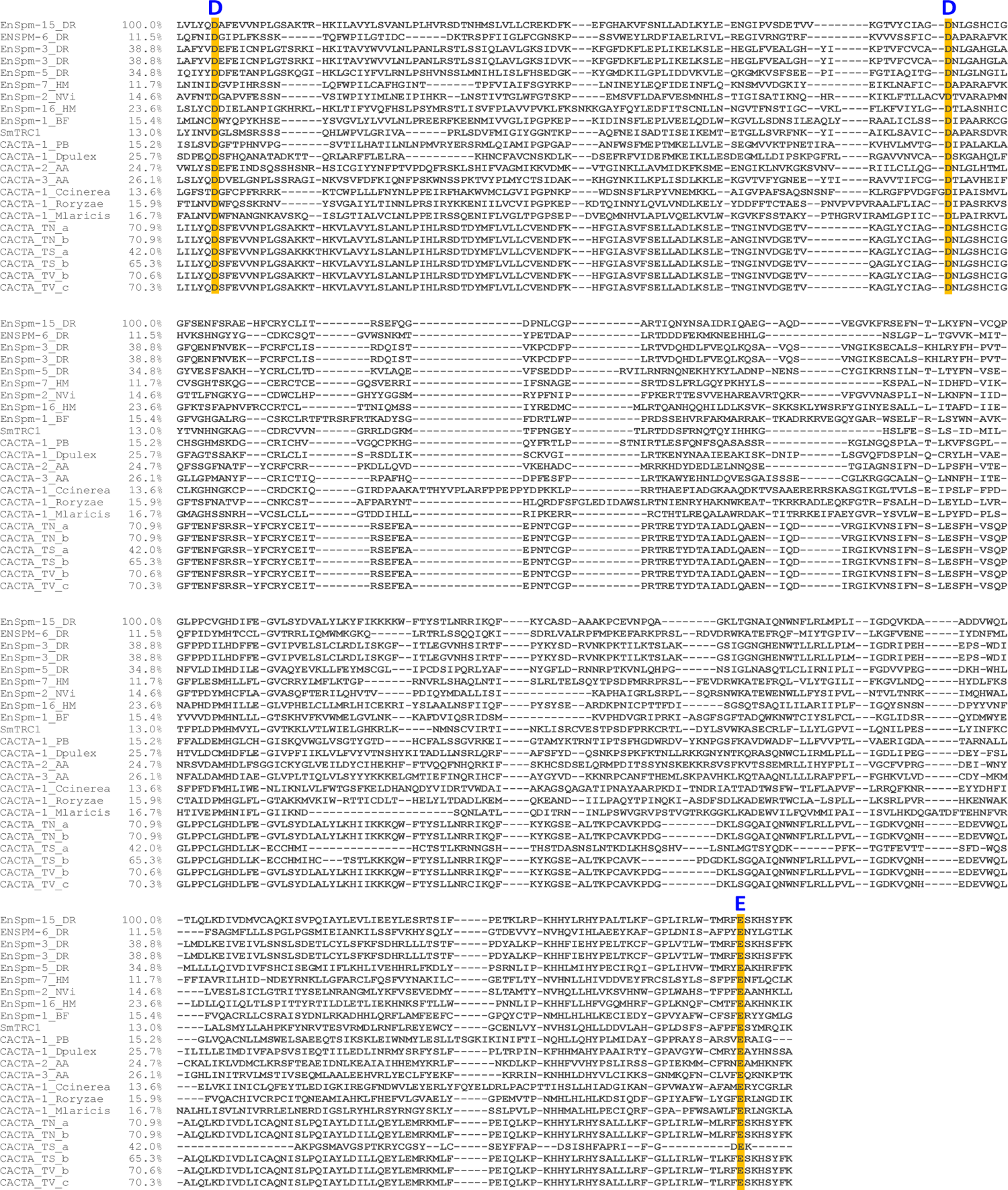
The catalytic center of the transposase contained in the CACTA transposon. The DDD/E triad is highlighted in orange in the protein sequence alignment and marked with blue letters according to (2, 3). CACTA_TN_a and CACTA_TN_b in *Takifugu niphobles* are 70% similar to the EnSpm-15_DR in zebrafish in terms of the protein-coding sequences residing within the DDD/E triad. Sequences were aligned by MUSCLE (https://www.ebi.ac.uk/Tools/msa/muscle/) and visualized in Mview (https://www.ebi.ac.uk/Tools/msa/mview/). AA, BF, DR, HM, NVi, PB, TN, TS, and TV represent *Aedes aegypti*, *Branchiostoma floridae*, *Danio rerio*, *Hydra vulgaris*, *Nasonia vitripennis*, *Phycomyces blakesleeanus*, *Takifugu niphobles*, *Takifugu snyderi*, and *Takifugu vermicularis*, respectively.

**Fig. S8.**
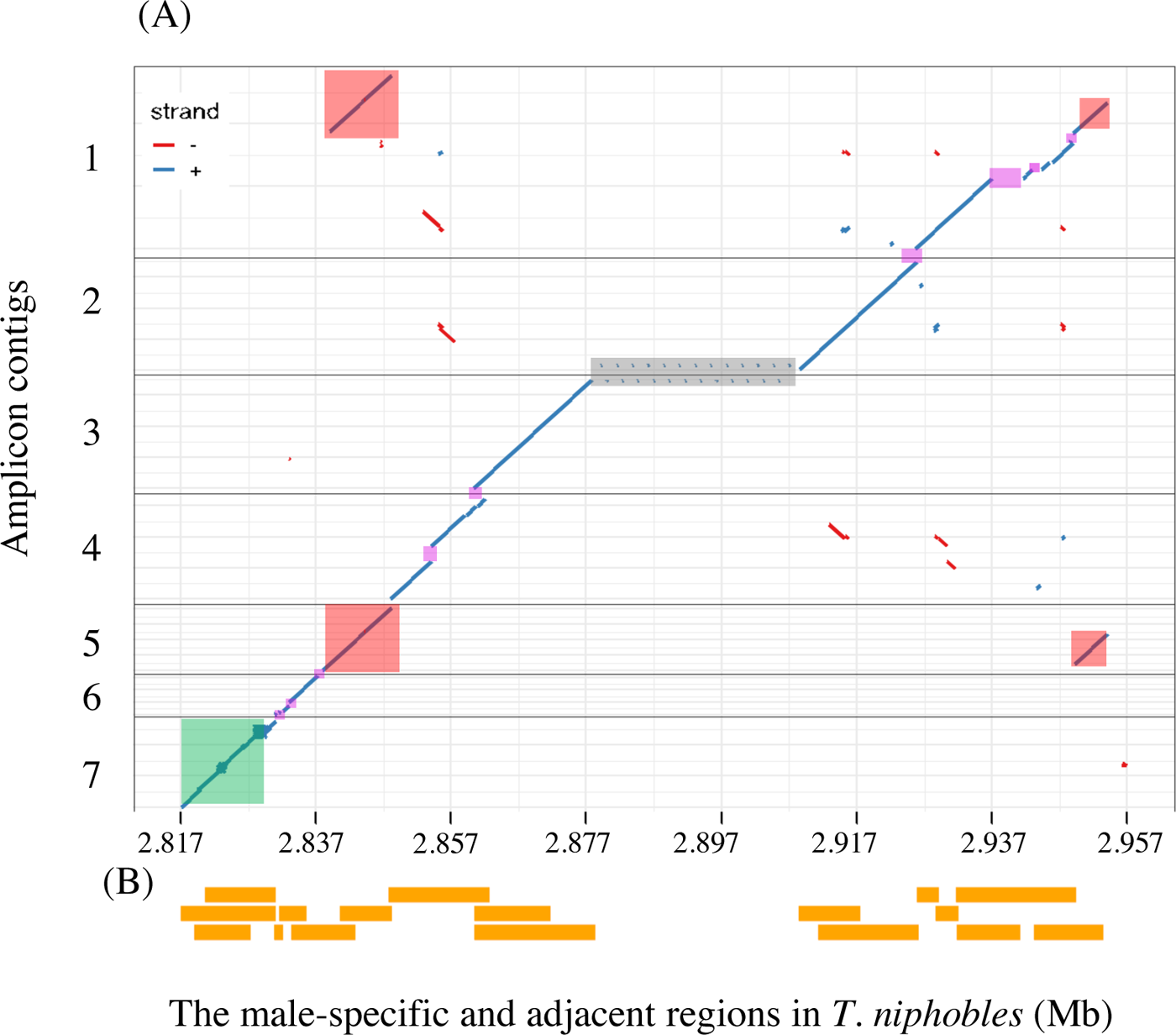
(A) Comparison between amplicon sequences and their targeted regions spanning the boundary between the pseudoautosomal region and the male-specific region in *T. niphobles*. Amplicon contigs were assembled using the sequences of tiled PCR products determined by Oxford Nanopore sequencing technology. No amplicons were generated in the grey shadow region due to the presence of ∼30-kb repetitive sequence at the 3’ end of *GsdfY* gene. Disconcordance between amplicon contigs and corresponding sequences on the *T. niphobles* Chr19Y assembly are shown in magenta shadows. Pink shadows indicate duplicated sequences in the male-specific region predicted from the Chr19Y assembly. The green shadow represents the pseudoautosomal region. (B) The predicted lengths of the PCR amplicons based on the Chr19Y assembly.

**Fig. S9.**
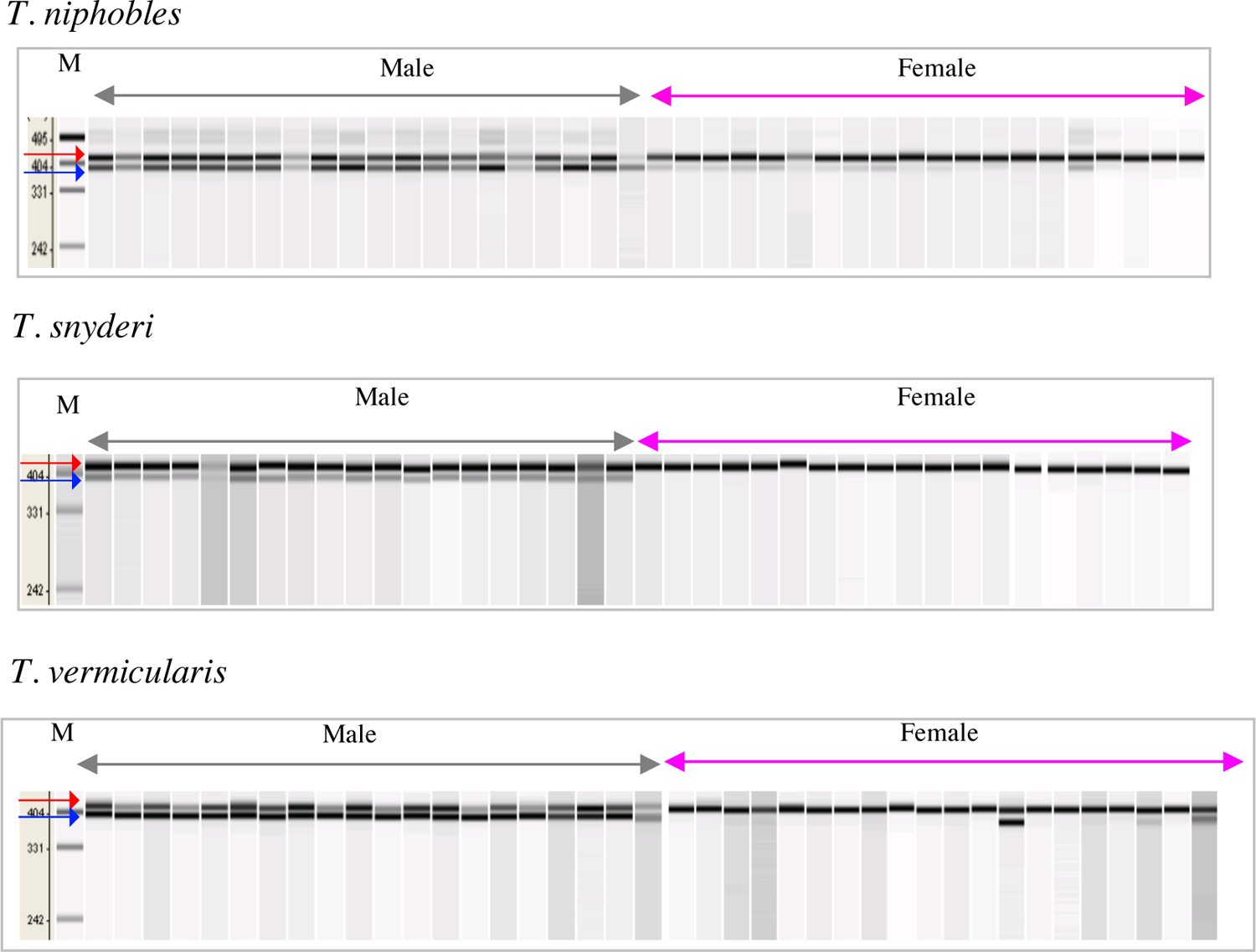
The presence of the male-specific region in wild populations of *T. niphobles*, *T. snyderi*, and *T. vermicularis*. Primer pairs that can produce paralog-specific amplicons (single-ended arrows) were used. Genotyping was performed in wild-caught individuals of each species. Amplified PCR products were analyzed on a MultiNA instrument. The lower band is associated with the male phenotype in each species. “M” denotes a set of molecular weight markers.

**Fig. S10.**
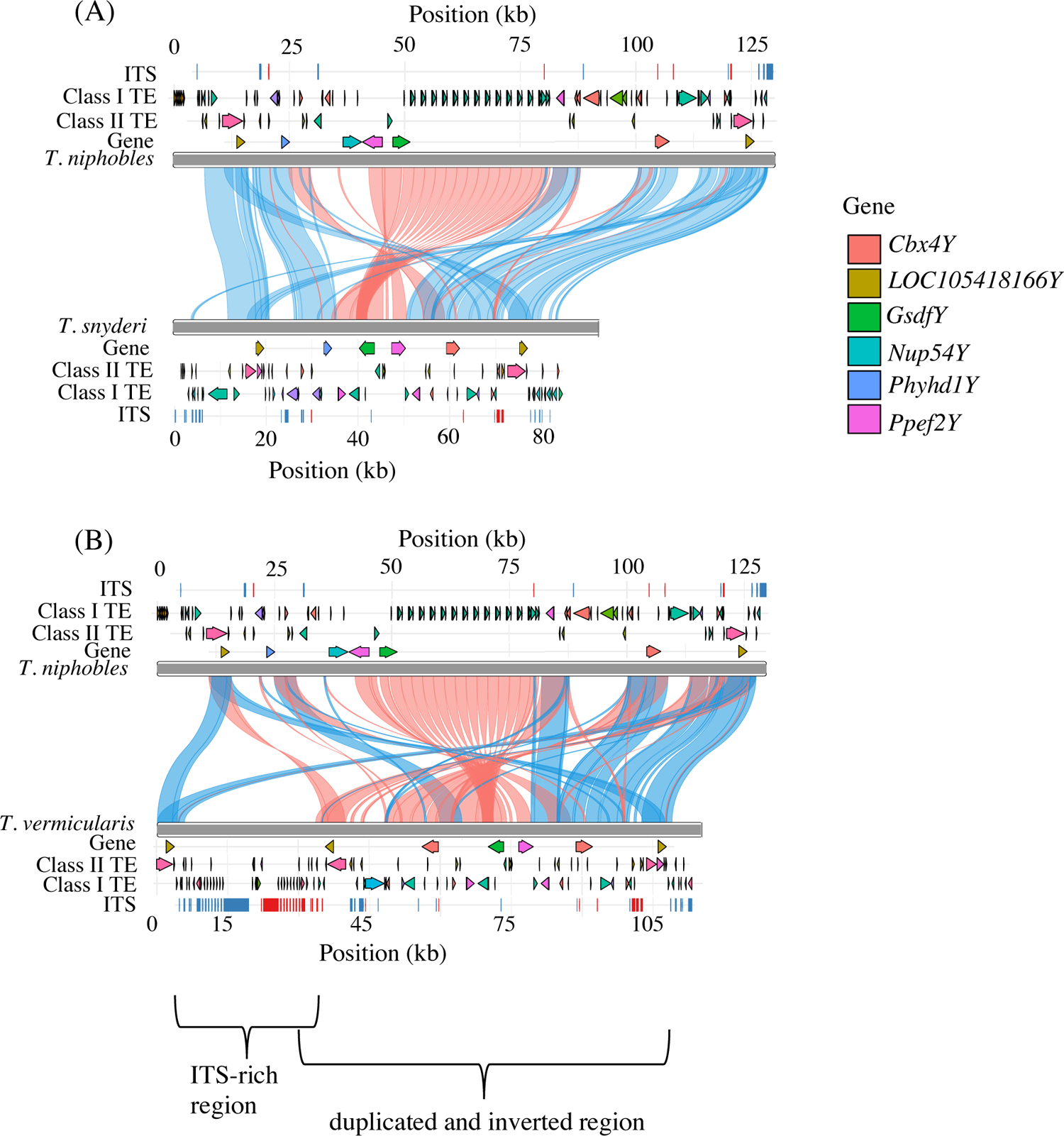
Schematic representations of the repeat annotation in the male-specific region in *T. niphobles*, *T. snyderi*, and *T. vermicularis*. Arrows depict Class I transposable elements (TEs), Class II TEs, and full-length genes. Rectangles represent interstitial telomeric sequences (ITSs) (TTAGGG)_n_ Syntenic and inverted segments are connected by blue and red ribbons, respectively. (A) Comparison of the male-specific region between *T. niphobles* and *T. snyderi*. (B) Comparison of the male-specific region between *T. niphobles* and *T. vermicularis*.

**Fig. S11.**
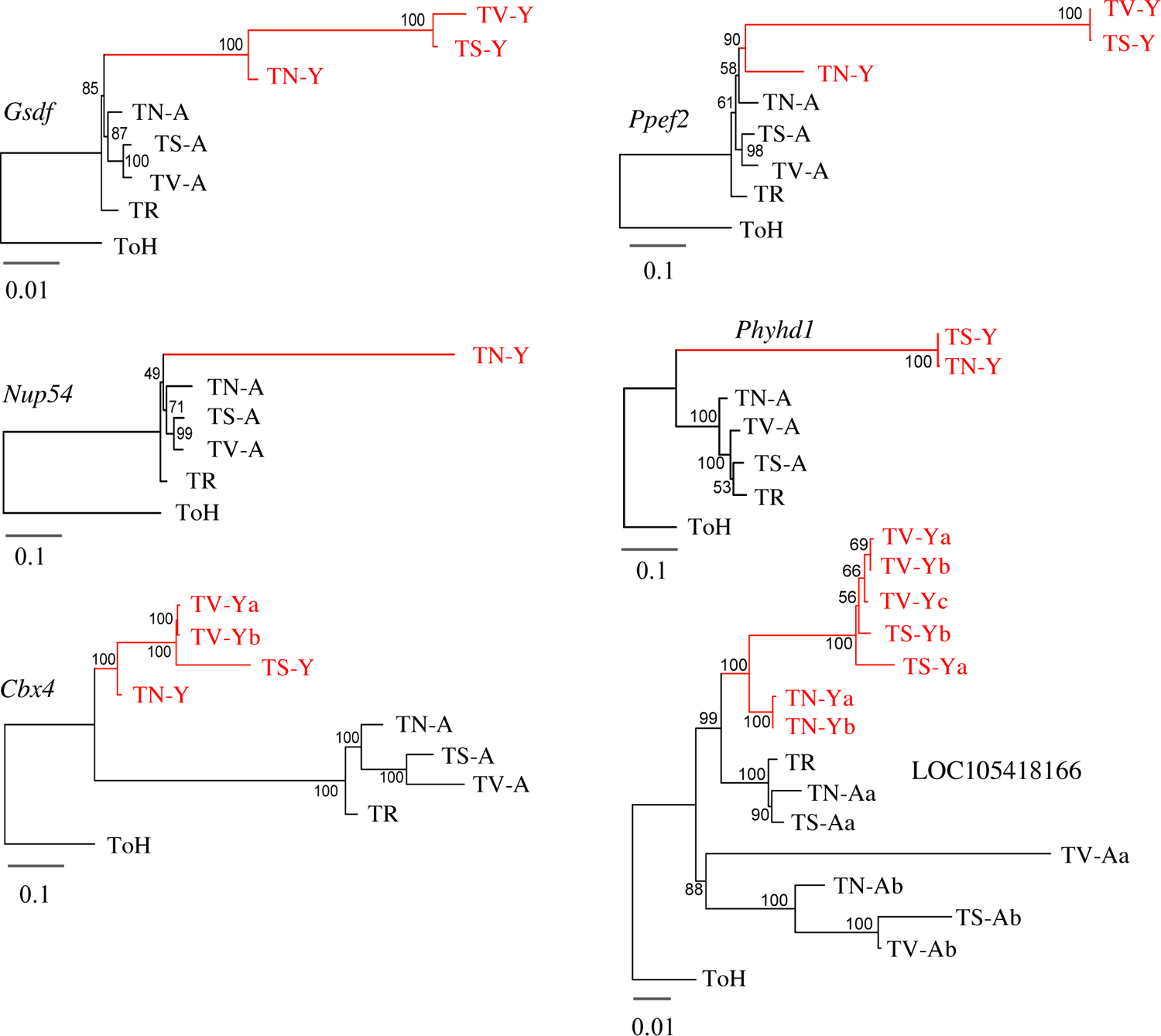
Maximum-likelihood clustering of the male-specific and autosomal paralogs in *Takifugu niphobles*, *T. snyderi*, and *T. vermicularis* incorporating InDels information. To reduce the possible effects of long-branch attraction, gaps were treated as fifth states in the multiple sequence alignment. A maximum-likelihood tree was built for each of the genes using RAxML (v. 8.0) with the MULTICAT model, GTR substitution model, and -V option. TR, TN, TS, and TV represent *T. rubripes*, *T. niphobles*, *T. snyderi*, and *T. vermicularis*, respectively. *Torquigener hypselogeneion* (ToH) sequences were used as the outgroup. The reliability of the inferred tree was tested by 1,000 fast bootstrap replicates. Red and black colors represent the male-specific (-Y) and autosomal (-A) sequences, respectively.

**Fig. S12.**
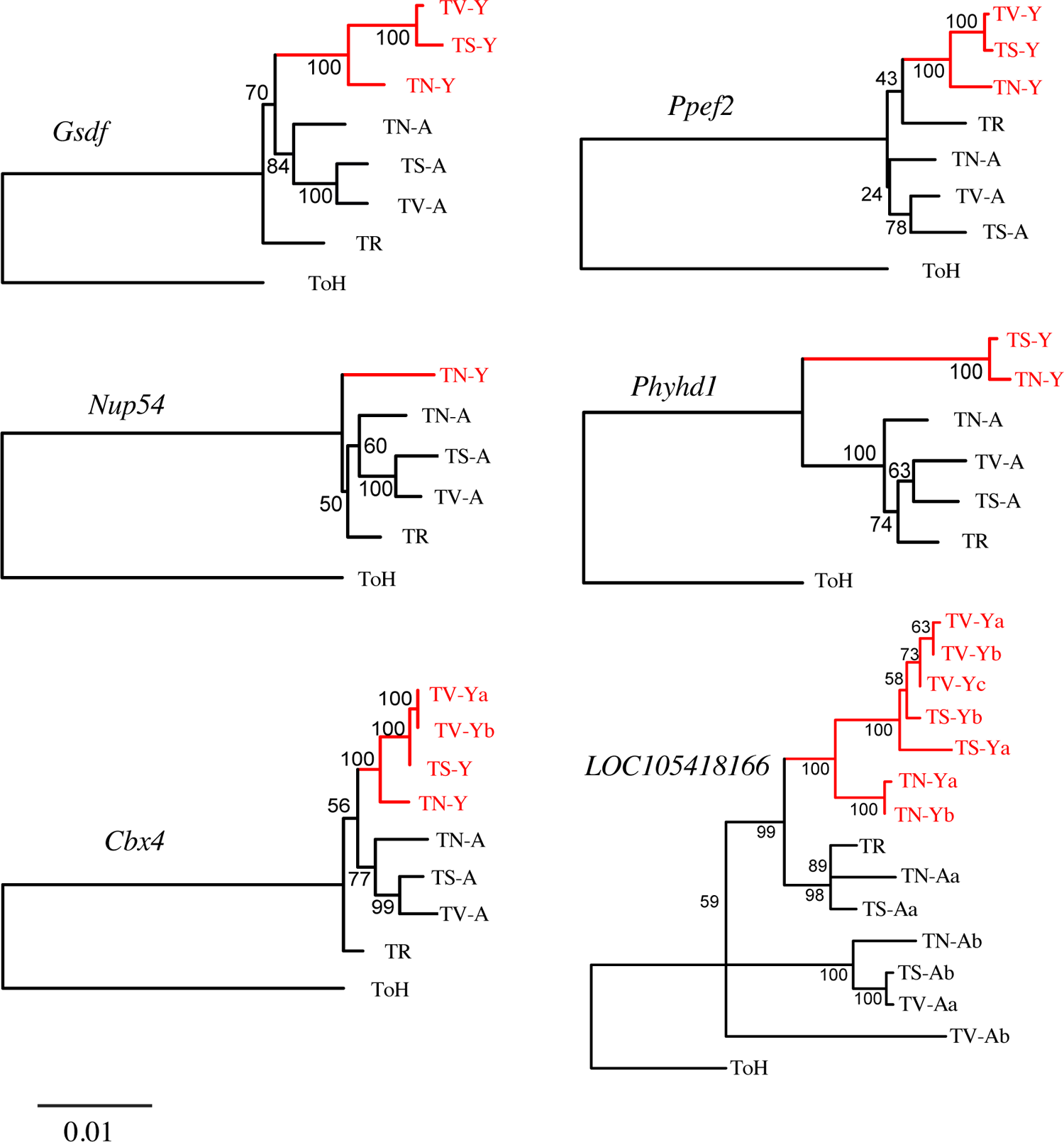
Maximum-likelihood clustering of the male-specific and autosomal paralogs in *Takifugu niphobles*, *T. snyderi*, and *T. vermicularis*. In contrast to Fig. S11, gap information was not incorporated in phylogenies. A maximum-likelihood tree was built for each of the genes using RAxML (v. 8.0) with the *GTRGAMMA* model and *--JC69* option. TR, TN, TS, and TV represent *T. rubripes*, *T. niphobles*, *T. snyderi*, and *T. vermicularis*, respectively. *Torquigener hypselogeneion* (ToH) sequences were used as the outgroup. The reliability of the inferred tree was tested by 1,000 fast bootstrap replicates. Red and black colors represent the male-specific (-Y) and autosomal (-A) sequences, respectively.

**Fig. S13.**
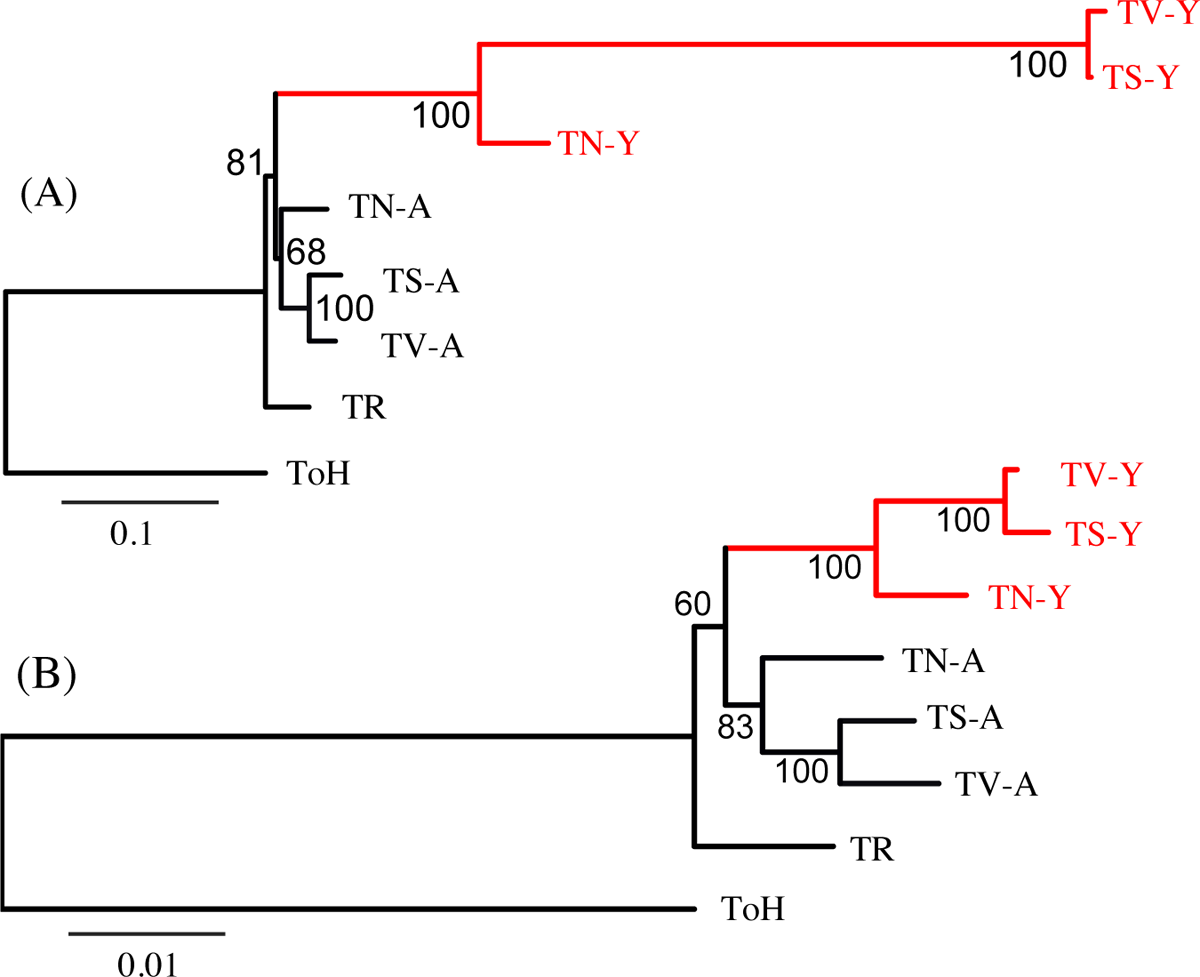
Maximum-likelihood clustering for the concatenated sequence of the male-specific and autosomal *Gsdf* and *Ppef2* genes in *Takifugu niphobles*, *T. snyderi*, and *T. vermicularis*. Since segmental duplication encompassing *Gsdf*, *Ppef2,* and *Nup54* on Chr6 likely contributed to the formation of the male-specific sequence in an ancestor of the three species, we concatenated *Gsdf* and *Ppef2* sequences and constructed phylogenetic trees. Note that *Nup54* was excluded since the male-specific paralog of this gene is absent in *T. snyderi* and *T. vermicularis*. (A) The tree incorporating InDels information. We treated gaps as fifth states in the multiple sequence alignment to incorporate InDels information. The tree was built using RAxML (v. 8.0) with the *MULTICAT* model and *GTR* substitution model. (B) The tree without incorporation of the InDels information. The tree was built using RAxML (v. 8.0) with the *GTRGAMMA* model. TR, TN, TS, and TV represent *T. rubripes*, *T. niphobles*, *T. snyderi*, and *T. vermicularis*, respectively. *Torquigener hypselogeneion* (ToH) sequences were used as the outgroup. The reliability of the inferred tree was tested by 1,000 fast bootstrap replicates. Red and black colors represent the male-specific (-Y) and autosomal (-A) sequences, respectively.

**Fig. S14.**
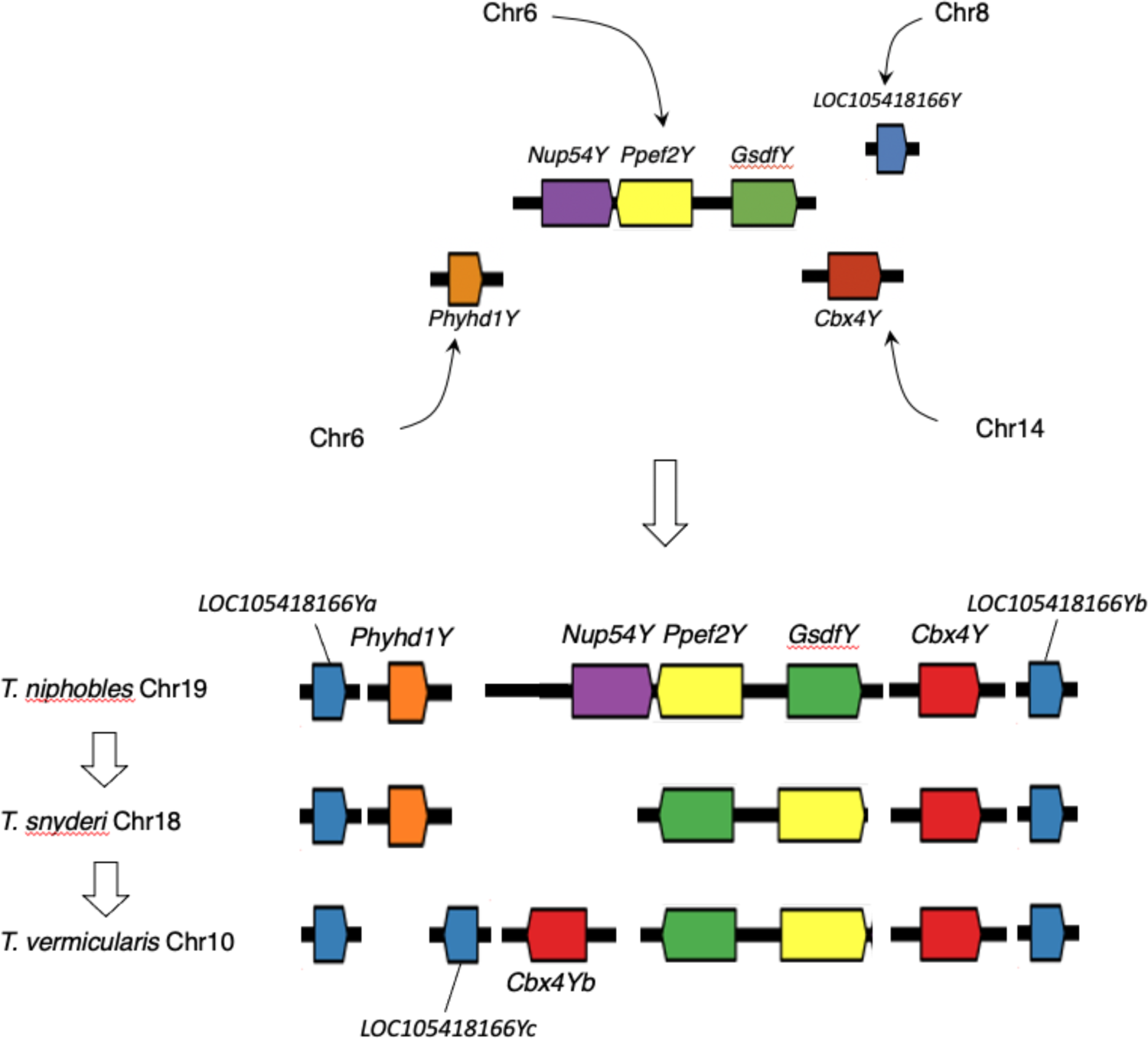
A hypothetical model for the evolutionary development of the core male-specific region. It is likely that the core male-specific region evolved in the common ancestor of the three species through the combination of at least two processes. One includes segmental duplication of the region encompassing *Nup54*, *Ppef2*, and *Gsdf* and the translocation of this region to the future sex chromosome. The other process is the gathering of other unlinked genes (*Phyhd1*, *Cbx4*, and *LOC105418166*) through independent duplications and translocations. *Nup54Y* was likely lost before the divergence of *T. snyderi* and *T. vermicularis*, while the gene content was retained in the lineage leading to *T. niphobles*. Then, in the *T. vermicularis* lineage, while *Phyhd1Y* was lost, the segmental duplication of part of the male-specific region resulted in the emergence of *LOC105418166Yc* and *Cbx4Yb*.

